# Molecular conflicts disrupting centromere assembly contribute to *Xenopus* hybrid inviability

**DOI:** 10.1101/2022.03.06.483208

**Authors:** Maiko Kitaoka, Owen K. Smith, Aaron F. Straight, Rebecca Heald

**Affiliations:** Department of Molecular and Cell Biology, University of California, Berkeley, CA 94720-3200, USA; Department of Biochemistry, Stanford University School of Medicine, Stanford, CA 94305-5307, USA

**Keywords:** Centromere, hybrid incompatibility, speciation, *Xenopus*, chromosome segregation, CENP-A

## Abstract

Although central to evolution, the causes of hybrid inviability that drive reproductive isolation are poorly understood. Embryonic lethality occurs when eggs of the frog *X. tropicalis* are fertilized with either *X. laevis* or *X. borealis* sperm. We observed that distinct subsets of paternal chromosomes failed to assemble functional centromeres, causing their mis-segregation during embryonic cell divisions. Core centromere DNA sequence analysis revealed little conservation among the three species, indicating that epigenetic mechanisms that normally operate to maintain centromere integrity are disrupted on specific paternal chromosomes in hybrids. In vitro reactions combining *X. tropicalis* egg extract with either *X. laevis* or *X. borealis* sperm chromosomes revealed that paternally matched or over-expressed centromeric histone CENP-A and its chaperone HJURP could rescue centromere assembly on affected chromosomes in interphase nuclei. However, whereas the *X. laevis* chromosomes maintained centromeric CENP-A in metaphase, *X. borealis* chromosomes did not, and also displayed ultra-thin regions containing ribosomal DNA. Both centromere assembly and morphology of *X. borealis* mitotic chromosomes could be rescued by inhibiting RNA Polymerase I or by preventing collapse of stalled DNA replication forks. These results indicate that specific paternal centromeres are inactivated in hybrids due to disruption of associated chromatin regions that interfere with CENP-A incorporation, at least in some cases due to conflicts between replication and transcription machineries. Thus, our findings highlight the dynamic nature of centromere maintenance and its susceptibility to disruption in vertebrate interspecies hybrids.

**ONE SENTENCE SUMMARY:** Centromere incompatibilities in inviable *Xenopus* hybrids are sequence-independent and result from disruption of epigenetic pathways required for centromere maintenance.

## INTRODUCTION

Hybridization between closely related species often leads to embryonic lethality accompanied by defects in genome stability and maintenance, but the cellular and molecular mechanisms underlying post-zygotic barriers that drive reproductive isolation and speciation are largely unknown (Sanei et al., 2011; Maheshwari and Barbash, 2011; Fujiwara et al., 1997; Gernand et al., 2005). Among animals, a number of studies of inviable hybrids resulting from crosses of related *Drosophila* species have revealed an important role for the centromere, the chromosomal site where the kinetochore assembles to mediate chromosome attachment to the mitotic spindle and segregation to daughter cells. Both centromere DNA sequence and protein components including the centromeric histone H3 variant, Centromere Protein A (CENP-A) are rapidly evolving (Malik and Henikoff, 2001; Maheshwari et al., 2015). Localization of exogenously expressed CENP-A to centromeres across *Drosophila* species was shown to require co-expression of its species-matched chaperone CAL1/HJURP, indicating that the CENP-A deposition machinery also co-evolves (Rosin and Mellone, 2016). In turn, kinetochore formation at centromeres depends on specific, epigenetic recognition and stabilization of CENP-A nucleosomes by other factors, including CENP-C, CENP-N, and M18BP1 (Moree et al., 2011; Carroll et al., 2009; Pentakota et al., 2017; Chittori et al., 2018; Tian et al., 2018; Falk et al., 2015; French et al., 2017; Shono et al., 2015; Hori et al., 2017). Thus, co-evolution of centromere DNA and many associated proteins generate barriers to hybrid viability by interfering with assembly of the chromosome segregation machinery.

Increasing evidence suggests that the chromatin environment also plays an important role in centromere assembly and that changes in the nuclear organization are related to hybridization outcomes. For example, disruption of the chromocenter, a domain containing the pericentromeric satellite DNA, is common among *Drosophila* hybrids and may underlie inviability (Jagannathan and Yamashita, 2021). Furthermore, known inviability factors such as hybrid male rescue (Hmr) and lethal hybrid rescue (Lhr) strongly impact chromosome segregation in *Drosophila* hybrids and have been reported to regulate transposable elements and heterochromatic repeats (Thomae et al., 2013; Satyaki et al., 2014), associate with chromatin chaperones adjacent to centromeres (Anselm et al., 2018), and to link pericentromeric and centromeric chromatin to maintain centromere integrity (Lukacs et al., 2021). However, whether these factors play a direct role in centromere function is unclear (Blum et al., 2017). Despite these advances, the relative contribution to hybrid inviability of diverging centromere sequences versus the activity and spatial organization of associated chromatin machineries that promote centromere assembly is poorly understood.

Among vertebrates, hybridization resulting in post-zygotic death has been more difficult to study. *Xenopus* frog species possess interesting evolutionary relationships that include past interspecies hybridization events (Session et al., 2016) and provide an ideal system to study the molecular basis of hybridization outcomes, since cross fertilization experiments are easily performed (Narbonne et al., 2011; Gibeaux et al., 2018), and mechanisms underlying hybrid incompatibility can be uniquely and powerfully investigated *in vitro* by combining the sperm chromosomes and egg extracts from different species. We showed previously that interspecies hybrids produced when *X. laevis* or *X. borealis* eggs are fertilized by *X. tropicalis* sperm are viable, while the reverse crosses die before gastrulation and zygotic gene activation by explosive cell lysis or exogastrulation, respectively (Gibeaux et al., 2018). The inviable hybrids displayed chromosome segregation defects during embryonic cleavages, characterized by lagging chromosomes, chromosome bridges, and formation of micronuclei. By whole genome sequencing, specific and distinct paternal chromosome regions were lost from both hybrids prior to embryo death. A fraction of *X. laevis* chromosomes failed to assemble centromeres/kinetochores, likely leading to spindle attachment defects and ultimately chromosome mis-segregation and embryo death (Gibeaux et al., 2018).

To better understand centromere-based *Xenopus* hybrid incompatibilities, here we combine genomic, in vitro, and in vivo analyses. We find that although core centromeric sequences are not conserved, *X. tropicalis* egg cytoplasm supports centromere assembly on *X. laevis* and *X. borealis* chromosomes. However, upon entry into metaphase, conflicts emerge that evict CENP-A from a subset of chromosomes. In the case of *X. laevis*, excess CENP-A and its chaperone HJURP can rescue this defect. In contrast, eviction of CENP-A from *X. borealis* chromosomes could be rescued by dissociating the rRNA polymerase Pol I or by preventing collapse of DNA replication forks. These results indicate that centromere incompatibility is driven primarily by centromere sequence-independent replication-transcription conflicts that disrupt the epigenetic maintenance of CENP-A nucleosomes.

## RESULTS

### Core centromere sequence variation does not underlie *Xenopus* hybrid aneuploidy

We previously observed chromosome mis-segregation and loss of centromere and kinetochore proteins from a subset of chromosomes in hybrids generated by fertilizing *X. tropicalis* eggs with *X. laevis* sperm. Whole genome sequencing just prior to embryo death revealed consistent deletion of large genomic regions from two paternal chromosomes, 3L and 4L (Gibeaux et al., 2018). We hypothesized that chromosome-specific aneuploidy resulted from divergent centromeric sequences on the affected chromosomes, rendering them incompatible with the maternal *X. tropicalis* centromeric histone CENP-A and its loading machinery. Recent characterization of *X. laevis* centromere sequences by chromatin immunoprecipitation with CENP-A antibodies and sequencing analysis (ChIP-seq) revealed a family of related sequences found in distinct combinations and abundances on different *X. laevis* chromosomes (Smith et al., 2021). However, *X. laevis* centromeres 3L and 4L did not possess any distinguishing features in terms of size or composition. Thus, differences in core centromere DNA sequences do not appear to drive the specific chromosome mis-segregation events and genome loss observed in the inviable *X. tropicalis*/*X. laevis* hybrid.

To expand our analysis, we characterized a second inviable hybrid resulting from fertilization of *X. tropicalis* eggs with sperm from *X. borealis*, a frog species possessing an allotetraploid genome closely related to *X. laevis* (Session et al., 2016). These hybrids display specific and consistent genome loss from a different subset of paternal chromosomes including 1S, 5S, 4L, and 8L (Gibeaux et al., 2018). To determine the extent to which centromere sequences differed across the three *Xenopus* species, CENP-A ChIP-seq was similarly applied to *X. tropicalis* and *X. borealis*. We used an alignment-independent *k*-mer based analysis to identify sequence features of the highly repetitive centromeric arrays in each species without the need for a complete genome sequence (Fig. S1A). Comparing the enrichment value (normalized CENP-A *k*-mer counts/normalized input *k*-mer counts) revealed that the majority of individual *k*-mers are enriched in one species, but not the others (Fig. 1A). Furthermore, analysis of full-length sequencing reads that contained CENP-A enriched *k*-mers showed that CENP-A nucleosome-associated DNA sequences of the three species bear little relationship to one another (Fig. 1B). These findings reinforce the idea that incompatibilities leading to mis-segregation of specific chromosomes are not due to centromere sequence differences. Interestingly, although protein sequence alignments of *X. laevis*, *X. tropicalis*, and *X. borealis* CENP-A showed that they are nearly 90% identical, divergence occurred in both the N-terminus and the CENP-A Targeting Domain (CATD) L1 loop region that provides specificity for recognition of the CENP-A/H4 complex by its dedicated chaperone HJURP (Fig. S1B) (Rosin and Mellone, 2016; Foltz et al., 2009; Dunleavy et al., 2009; Hu et al., 2011). Together, these results suggest that as in *Drosophila*, centromere sequences, CENP-A, and its chaperone have co-evolved in *Xenopus* to strengthen specificity of their interactions (Rosin and Mellone, 2017). However, although disruption of these interactions can lead to inviability in flies (Henikoff et al., 2011; Malik and Henikoff, 2009), differences among core centromere sequences and CENP-A proteins does not explain loss of centromere function on a subset of chromosomes in inviable *Xenopus* hybrids.

**FIGURE 1:**
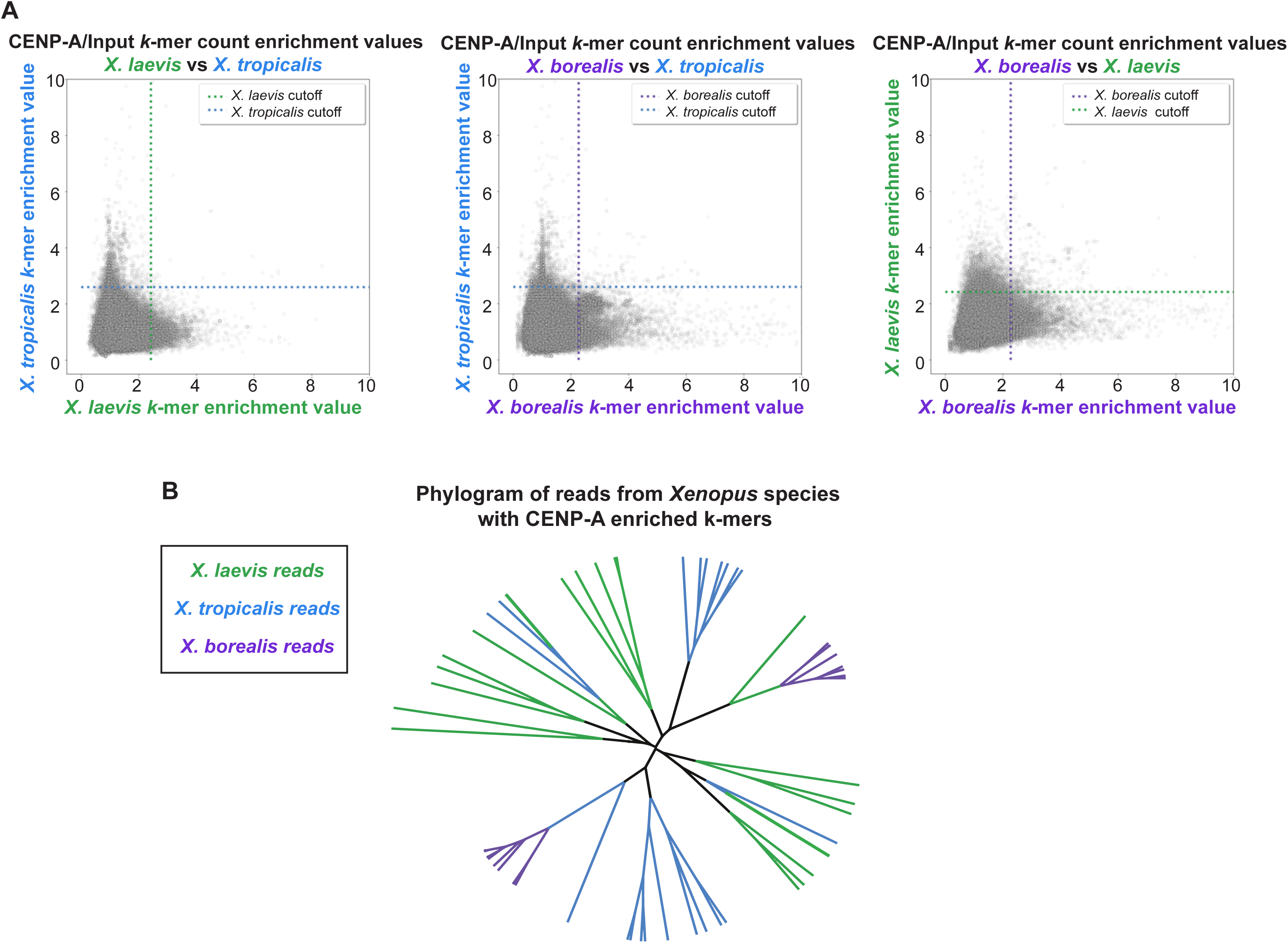
Comparison of *X. laevis*, *X. tropicalis*, and *X. borealis* core centromere sequences. (**A**) Scatter plots of *k*-mer enrichment values (normalized CENP-A counts / normalized input counts) compared between species. Only *k*-mers found in both species are plotted. Dotted lines indicate enrichment value for each species that is five median absolute deviations above the median enrichment value to denote highly enriched *k*-mers, which are not well conserved across species. (**B**) Phylogram of full-length sequencing reads from each *Xenopus* species. Branches are colored according to species of origin. Sequencing reads were selected first by the presence of at least 80 CENP-A enriched 25bp *k*-mers and then by hierarchical clustering. The phylogram illustrates a striking divergence of core centromere sequences.

### CENP-A eviction from a subset of chromosomes requires cell cycle progression

To better understand the process by which specific chromosomes lose centromere function in hybrids, we took advantage of the *Xenopus* egg extract system capable of recapitulating events of the cell division cycle in vitro, including sperm chromosome replication, mitotic chromosome condensation, and centromere/kinetochore formation (Maresca and Heald, 2006; French and Straight, 2017). To monitor centromere assembly, *X. tropicalis, X. laevis*, or *X. borealis* sperm nuclei were added to *X. tropicalis* egg extracts and probed for CENP-A at different stages of the cell cycle. Sperm chromosomes of all three species condensed and possessed single centromeric CENP-A foci when added directly to metaphase-arrested *X. tropicalis* extract (Fig. 2A, B), consistent with observations that sperm chromosomes contain CENP-A (Bernad et al., 2011; Milks et al., 2009). However, cycling the extract through interphase to allow sperm decondensation, nuclear envelope formation, and DNA replication in *X. tropicalis* egg cytoplasm resulted in no visible CENP-A on a subset of *X. laevis* and *X. borealis* mitotic chromosomes in the subsequent metaphase, whereas *X. tropicalis* centromeres were not affected (Fig. 2A, B). To determine when in the cell cycle CENP-A was evicted from paternal chromosomes, we examined interphase nuclei in control and hybrid in vitro reactions. The expected number of centromere foci, 18 for *X. laevis* and *X. borealis*, decreased in *X. tropicalis* extract (Fig. 2C, Fig S2A, B). The loss of CENP-A localization from 2 or 4 paternal *X. laevis* and *X. borealis* chromosomes, respectively, corresponded very well to whole genome sequencing data of hybrid embryos in terms of the number of chromosomes affected (Gibeaux et al., 2018), and indicates that CENP-A is lost from this subset of paternal chromosomes during interphase.

**FIGURE 2:**
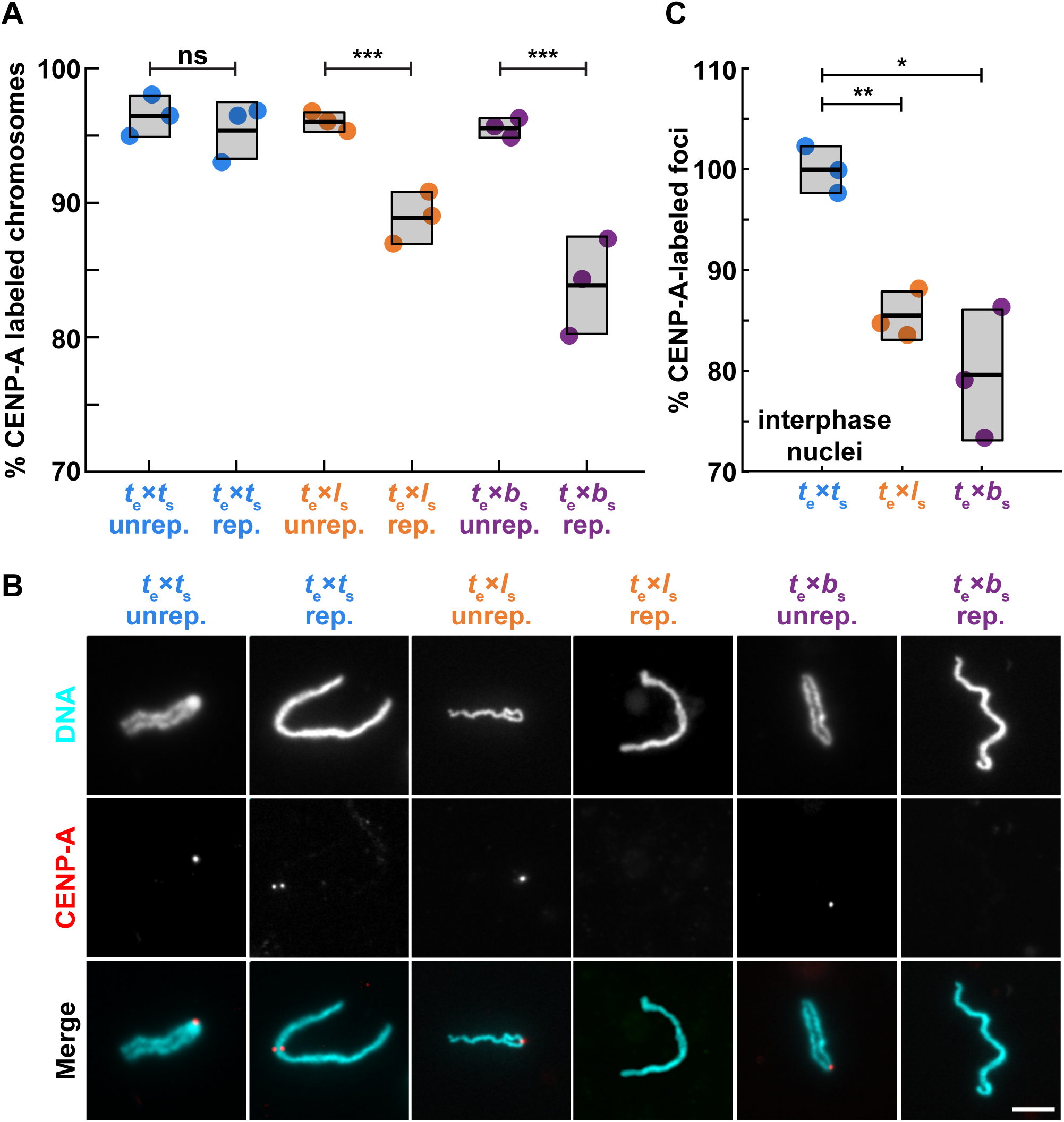
Loss of centromeric CENP-A is cell cycle-dependent. (**A**) Percentage of mitotic chromosomes with centromeric CENP-A staining in *X. tropicalis* egg extract. Over 95% of *X. tropicalis*, *X. laevis*, and *X. borealis* unreplicated sperm chromosomes added directly to metaphase-arrested *X. tropicalis* egg extracts possess centromeres, as indicated by immunofluorescence of the centromeric histone CENP-A. Following progression through the cell cycle, a fraction of replicated *X. laevis* and *X. borealis* mitotic chromosomes completely lose centromeric CENP-A foci. Unrep., unreplicated chromosomes; rep, replicated chromosomes. N = 3 extracts, N > 275 chromosomes per extract. p-values (left to right) by two-tailed two-sample unequal variance t-tests: 0.3356, 0.0008, 0.0004; ns, not significant. (**B**) Representative images of mitotic unreplicated and replicated *X. tropicalis*, *X. laevis*, and *X. borealis* chromosomes formed in *X. tropicalis* egg extracts. DNA in cyan, CENP-A in red. Scale bar is 10 µm. (**C**) Percentage of total expected CENP-A foci observed in nuclei formed in interphase *X. tropicalis* egg extract. *X. laevis* and *X. borealis* interphase nuclei both lose centromere foci during interphase, prior to entry into metaphase, whereas *X. tropicalis* nuclei do not. From N = 3 extracts, N > 64 nuclei per extract. p-values (top to bottom) by one-way ANOVA with Tukey post-hoc analysis: 0.0025, 0.0133. Species nomenclature throughout figures denotes egg extract as subscript e and chromosomes as subscript s, for example *t*_e_ x *l*_s_ indicates *X. tropicalis* egg extract combined with *X. laevis* sperm chromosomes. *X. tropicalis* is color-coded blue, while *X. laevis* and *X. borealis* hybrid combinations are orange and purple, respectively.

The recent detailed characterization of *X. laevis* centromere sequences allowed us to test whether the centromere assembly defects observed in egg extract occurred on the same chromosomes disrupted in hybrid embryos (Smith et al., 2021; Gibeaux et al., 2018). Fluorescence in situ hybridization (FISH) probes in combination with CENP-A immunofluorescence identified *X. laevis* chromosomes 3L and 4L, the two chromosomes that lose large genomic regions in hybrid embryos, as those that also lose centromeric CENP-A staining when replicated in *X. tropicalis* egg extract (Fig. S2C-E). Thus, the in vitro system reproduces incompatibilities likely to underlie chromosome mis-segregation and ultimately genome loss observed in vivo. These results show that while all paternal sperm chromosomes initially possess CENP-A at their centromeres, a subset evict CENP-A during interphase, indicating that epigenetic mechanisms enable hybrid centromere assembly despite evolutionary differences, but are disrupted on individual chromosomes.

### CENP-A and its chaperone HJURP can rescue *X. laevis* centromere assembly

We next sought to determine whether enhancing centromere assembly by adding species-matched paternal factors could prevent CENP-A eviction and centromere loss from specific *X. laevis* and *X. borealis* chromosomes formed in *X. tropicalis* extracts. In vitro reactions were supplemented with paternally-matched proteins expressed in reticulocyte lysate, including CENP-A and its dedicated chaperone HJURP, at the onset of interphase (Fig. S3A-C). Whereas adding *X. laevis* CENP-A resulted in a partial rescue, CENP-A plus HJURP increased the percentage of replicated *X. laevis* mitotic chromosomes with CENP-A foci to control levels (Fig. 3A). In contrast, no combination of *X. borealis* centromere factors tested, including CENP-A, HJURP, and CENP-C (Erhardt et al., 2008; Roure et al., 2019), restored CENP-A foci to replicated *X. borealis* mitotic chromosomes (Fig. 3B). Notably, however, examination of interphase nuclei in *X. borealis* sperm/*X. tropicalis* egg extract reactions prior to metaphase entry revealed that CENP-A localization was initially fully rescued, with the expected number of CENP-A-positive foci corresponding to the number of chromosomes (Fig. 3C). These results indicate that exogenous species-matched CENP-A can restore proper centromere formation on all chromosomes during interphase of for both *X. laevis* and *X. borealis*, but that CENP-A is not maintained on a subset of *X. borealis* chromosomes upon entry into mitosis.

**FIGURE 3:**
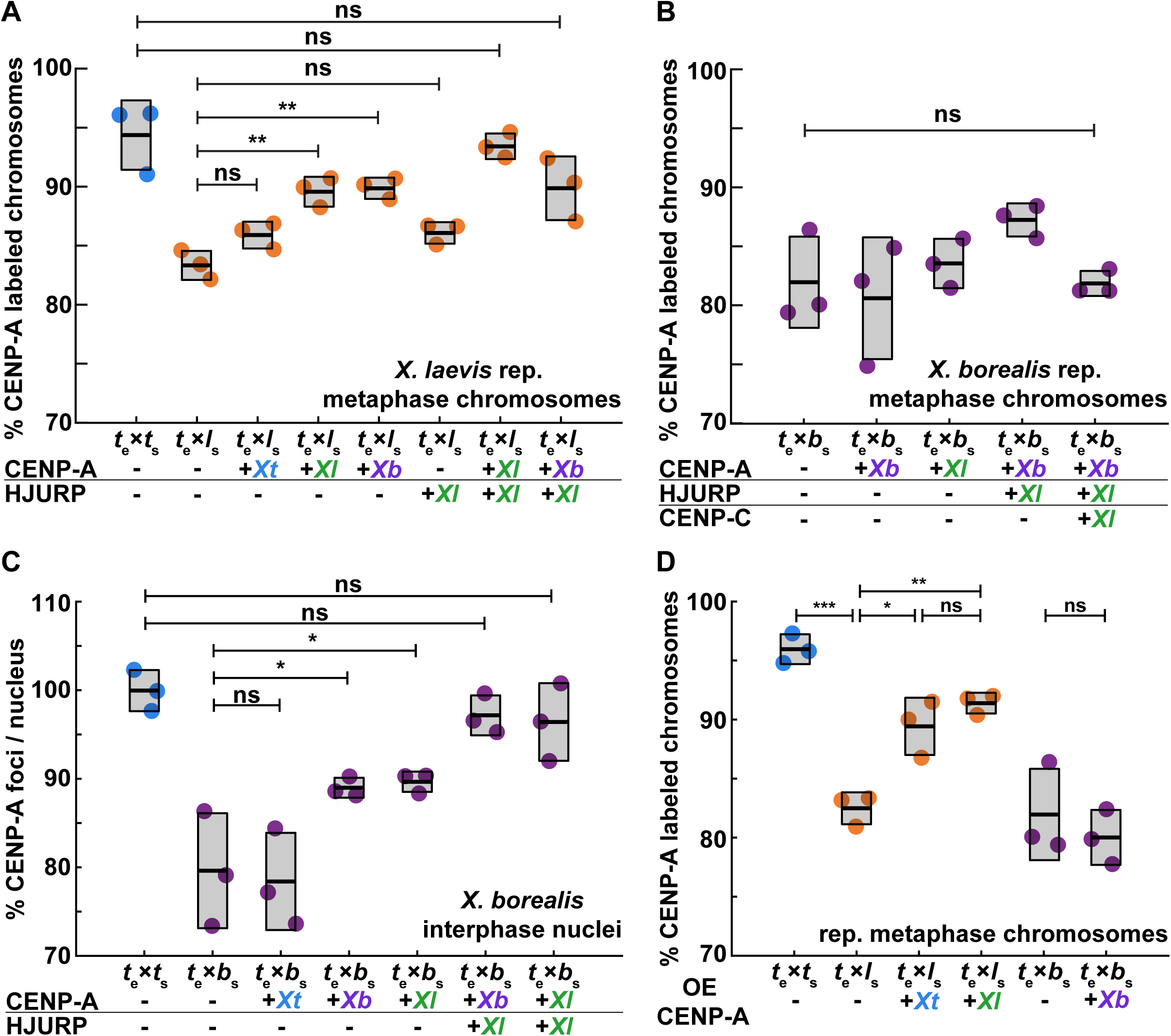
Driving CENP-A assembly rescues centromere localization in interphase, which persists on mitotic *X. laevis*, but not on *X. borealis* chromosomes. (**A**) Percentage of replicated *X. laevis* chromosomes with centromeric CENP-A staining in *X. tropicalis* extract supplemented with in vitro translated CENP-A and HJURP proteins from different *Xenopus* species. *X. laevis* chromosomes are fully rescued with species-matched centromere proteins. Quantification with N = 3 extracts, N > 315 chromosomes per extract. p-values (top to bottom) by one-way ANOVA with Tukey post-hoc analysis: 0.1734, 0.9999, 0.5522, 0.0057, 0.0086, 0.6281. (**B**) Percentage of replicated *X. borealis* chromosomes with centromeric CENP-A staining in *X. tropicalis* extract supplemented with in vitro translated centromere proteins from different *Xenopus* species. No combination or increased amounts of centromeric proteins CENP-A (CA), HJURP (HJ), and CENP-C (CC) restored CENP-A localization on *X. borealis* mitotic chromosomes. Quantification with N = 3 extracts, N > 216 chromosomes per extract. p-value by one-way ANOVA = 0.0786. (**C**) Percentage of CENP-A-labeled centromeric foci in *X. borealis* nuclei assembled in *X. tropicalis* extract supplemented with in vitro translated centromere proteins from different *Xenopus* species. Driving centromere assembly with species-matched proteins fully restores formation of centromere foci in interphase, but CENP-A staining is subsequently lost in metaphase (panel B). Quantification with N = 3 extracts, N > 67 nuclei per extract. p-values (top to bottom) by one-way ANOVA: 0.9996, 0.0562, 0.0433, 0.9690, 0.9109. (**D**) Percentage of replicated *X. laevis* or *X. borealis* chromosomes with centromeric CENP-A staining in *X. tropicalis* extract supplemented with excess (>50X endogenous levels) of in vitro translated *X. laevis* or *X. tropicalis* CENP-A. Whereas centromere staining is fully rescued on *X. laevis* mitotic chromosomes by CENP-A from either species, *X. borealis* centromere staining is not affected. Quantification with N = 3 extracts, N > 204 chromosomes per extract. p-values (top to bottom, then left to right) by one-way ANOVA with Tukey post-hoc analysis: 0.0042, 0.0001, 0.0249, 0.8845, 0.88946. A-D: ns, not significant.

The ability to mix and match egg extract, sperm chromosomes, and exogenous centromere assembly factors enabled evaluation of CENP-A/centromere compatibilities across species. For example, despite striking differences in core centromere sequences between *X. laevis* and *X. borealis* (Fig. 1), the CATDs of the two species’ CENP-A sequences are identical (Fig. S1B), and exogenous CENP-A from either species equivalently restored centromere assembly on *X. laevis* mitotic chromosomes replicated in *X. tropicalis* egg extract (Fig. 3A). Further, we observed that addition of excess exogenous *X. tropicalis* CENP-A could also increase the percentage of *X. laevis* mitotic chromosomes with centromere foci to control levels, although *X. borealis* chromosomes could not be rescued under any condition tested (Fig. 3B, D). Together, our results indicate that enhancing the pathway that drives CENP-A incorporation into centromeric chromatin can overcome whatever is destabilizing centromeres on specific *X. laevis* centromeres and raised the question of why the *X. borealis* chromosomes are refractory to this rescue.

### *X. borealis* chromosome defects result from mitotic replication stress

A clue as to why *X. borealis* mitotic chromosomes behave differently than *X. laevis* chromosomes in the in vitro hybrid extract system emerged with observation of their morphology. Although a subset of replicated *X. laevis* mitotic chromosomes formed in *X. tropicalis* extract lacked centromeres, they otherwise appeared normal. In contrast, 7-10% of *X. borealis* mitotic chromosomes displayed ultra-thin regions of 2-3 µm in length following replication, although centromeres on these chromosomes appeared largely intact (Fig. 4A, B; Fig. S4A, B). We reasoned that incomplete DNA replication leading to fork stalling and subsequent collapse in mitosis, termed replication stress, caused the formation of fragile sites (Gómez-González and Aguilera, 2019; Deng et al., 2019). Consistent with this idea, adding low doses of the DNA polymerase inhibitor aphidicolin that leads to replication stress (Deng et al., 2019; Kabeche et al., 2018; Durkin and Glover, 2007) triggered formation of ultra-thin regions on *X. tropicalis* and *X. laevis* mitotic sperm chromosomes that had progressed through the cell cycle in *X. tropicalis* extract, and slightly exacerbated morphological defects of *X. borealis* chromosomes (Fig. S4C-E). Notably, however, the aphidicolin-induced replication stress did not significantly affect CENP-A localization efficiency (Fig. S4F), indicating that replication stress per se does not interfere with CENP-A loading and maintenance.

**FIGURE 4:**
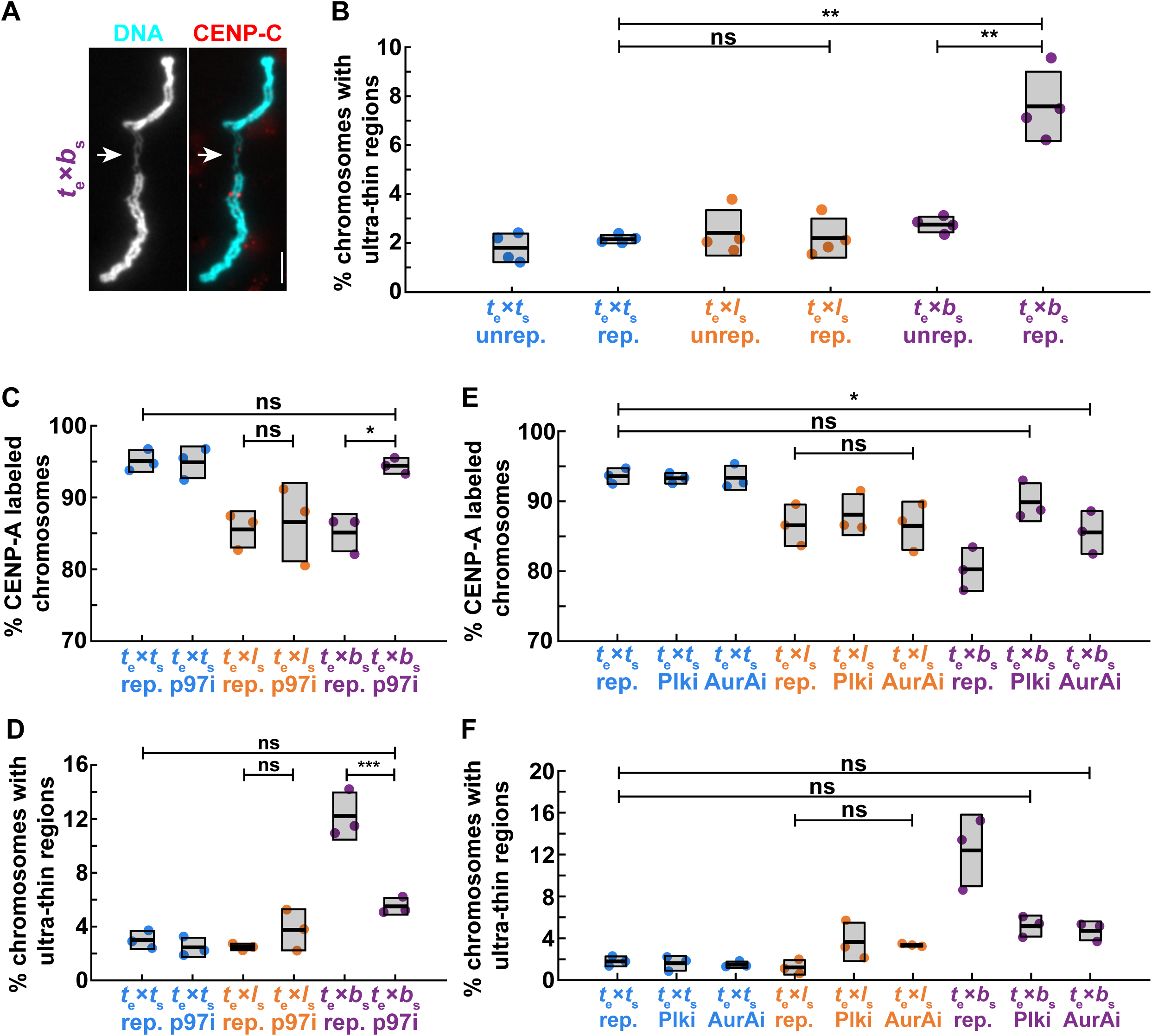
Mitotic replication stress leads to *X. borealis* centromere and chromosome morphology defects. (**A**) Representative image showing an ultra-thin region of a mitotic *X. borealis* chromosome formed in *X. tropicalis* egg extract. Note that the chromosome has an intact centromere. DNA in cyan, CENP-A in red. Scale bar is 5 µm. (**B**) Percentage of unreplicated and replicated mitotic chromosomes with ultrathin morphology defects in *X. tropicalis* extract. A low percentage of *X. tropicalis, X. laevis* or *X. borealis* unreplicated chromosomes display ultra-thin regions. After cycling through interphase, only *X. borealis* chromosomes exhibit a significant increase in this defect. Quantification with N = 3 extracts, N > 310 chromosomes per extract. p-values (top to bottom, then left to right) by one-way ANOVA with Tukey post-hoc analysis: 2.9352e-7, 0.9999, 1.6475e-6. (**C**) Percentage of replicated chromosomes with centromeric CENP-A staining in *X. tropicalis* extracts treated with solvent control or 10 µM p97 ATPase inhibitor NMS-873 (p97i). Inhibition of p97 restores CENP-A staining on *X. borealis* mitotic chromosomes, but does not affect *X. tropicalis* or *X. laevis* chromosomes. p-values (top to bottom, then left to right) by one-way ANOVA with Tukey post-hoc analysis: 0.9997, 0.9978, 0.0204. (**D**) Percentage of chromosomes with ultrathin regions in *X. tropicalis* extracts treated with solvent control or 10 µM p97 ATPase inhibitor NMS-873 (p97i). Inhibition of p97 rescues *X. borealis* chromosome morphology defects, but does not affect *X. tropicalis* or *X. laevis* chromosomes. p-values (top to bottom, then left to right) by one-way ANOVA with Tukey post-hoc analysis: 0.1114, 0.6903, 6.2572e-5. (**E**) Percentage of replicated chromosomes with centromeric CENP-A staining in *X. tropicalis* extracts treated with solvent control, 1 µM Polo-like kinase 1 inhibitor BI-2536 (Plk1i), or 1 µM Aurora A kinase inhibitor MLN-8237 (AurAi). CENP-A localization is fully or partially rescued on *X. borealis* mitotic chromosomes, whereas *X. tropicalis* or *X. laevis* chromosomes are not affected. p-values (top to bottom) by one-way ANOVA with Tukey post-hoc analysis: 0.0276, 0.7003, 0.9999. (**F**) Percentage of chromosomes with ultrathin regions in *X. tropicalis* extracts treated with solvent control, 1 µM Polo-like kinase 1 inhibitor BI-2536 (Plk1i), or 1 µM Aurora A kinase inhibitor MLN-8237 (AurAi). Inhibition of Plk1 and AurA rescued *X. borealis* mitotic chromosome morphology defects, but did not affect *X. tropicalis* or *X. laevis* chromosomes. p-values (top to bottom) by one-way ANOVA with Tukey post-hoc analysis: 0.2882, 0.1525, 0.5887. C, D: N = 3 extracts, N > 179 chromosomes per extract. E, F: N = 3 extracts, N > 155 chromosomes per extract. B-F: ns, not significant.

To determine whether the *X. borealis* chromosome morphology and mitotic centromere defects were due to replication stress, we blocked collapse of stalled mitotic replication forks by adding an inhibitor of the ATPase p97 (Fig. S4C), which is required for replication helicase removal (Maric et al., 2014; Deng et al., 2019). We observed a complete rescue of CENP-A localization and chromosome morphology on *X. borealis* chromosomes (Fig. 4C, D). Consistent with the factors known for this pathway of mitotic replication fork collapse and breakage (Deng et al., 2019), Aurora A and Plk1 kinase inhibitors added to *X. tropicalis* extracts at low doses that avoided mitotic defects also rescued *X. borealis* chromosome morphology and CENP-A localization, but did not affect *X. laevis* or *X. tropicalis* chromosomes (Fig. 4E, F). Finally, *X. laevis* or *X. tropicalis* chromosomes treated with aphidicolin followed by p97 inhibition displayed very few chromosome defects (Fig. S4C, D). Combined, these data reveal that a subset of *X. borealis* chromosomes experience mitotic replication stress in *X. tropicalis* cytoplasm, and that this is coupled to CENP-A eviction. Remarkably, however, centromere loss appears to occur on a different subset of mitotic chromosomes than those with ultra-thin regions.

### Replication-transcription conflicts lead to centromere defects

The fragile sites observed on *X. borealis* chromosomes were reminiscent of secondary constrictions that occur at repetitive, late-replicating regions such as ribosomal DNA (rDNA; Durica and Krider, 1977; Durkin and Glover, 2007). In *Xenopus*, the rDNA transcription machinery associates with mitotic chromosomes early in development and in egg extract (Roussel et al., 1996; Gébrane-Younès et al., 1997; Bell and Scheer, 1997; Bell et al., 1997), even though rDNA transcription and nucleolus formation occur after zygotic genome activation (Shiokawa et al., 1994; Newport and Kirschner, 1982). We therefore tested whether ultra-thin regions of *X. borealis* chromosomes replicated in *X. tropicalis* extract contained rDNA by performing immunofluorescence using antibodies against RNA polymerase I (Pol I) and the rDNA transcription regulator upstream binding factor (UBF). Both proteins were consistently enriched on the ultra-thin regions of *X. borealis* mitotic chromosomes assembled in *X. tropicalis* egg extract (Fig. 5A, B).

**Figure 5:**
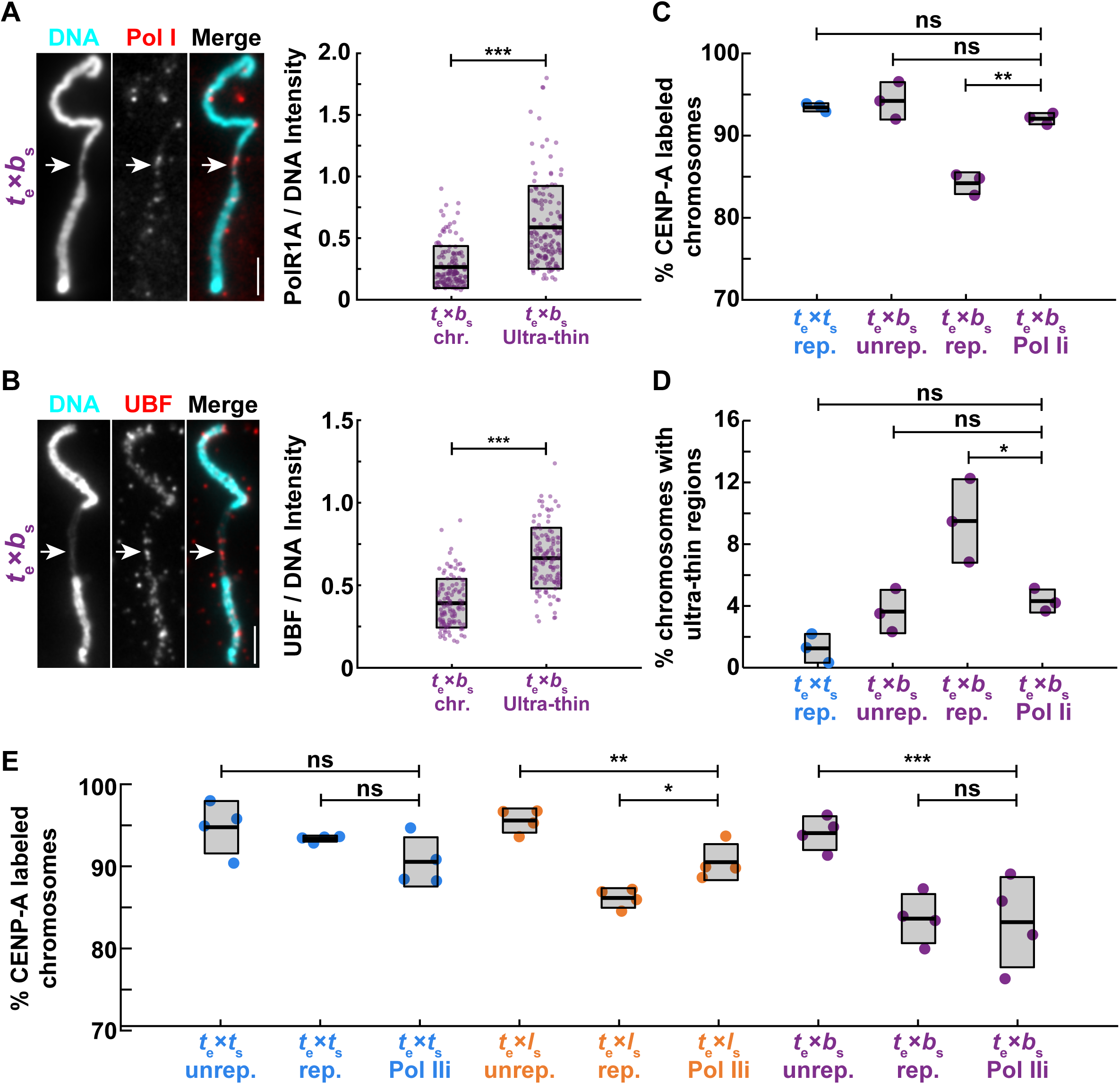
Replication-transcription conflicts at rDNA on *X. borealis* chromosomes can be rescued by inhibiting RNA Pol I. (**A**) Representative images and fluorescence intensity quantification of RNA Pol I staining relative to DNA on ultrathin and normal regions of *X. borealis* mitotic chromosomes, revealing enrichment of Pol I on ultra-thin regions. Quantification with N = 3 extracts, N = 140 chromosomes. p-value = 9.4793e-20 by two-tailed two-sample unequal variance t-tests. (**B**) Representative images and fluorescence intensity quantification of UBF staining relative to DNA on ultrathin and normal regions of *X. borealis* mitotic chromosomes, revealing enrichment of UBF on ultra-thin regions. Quantification with N = 3 extracts, N = 62 chromosomes. p-value = 4.5004e-13 by two-tailed two-sample unequal variance t-tests. (**C**) Percentage of mitotic chromosomes with centromeric CENP-A staining in *X. tropicalis* extracts treated with solvent control or 1 µM BMH-21 to inhibit RNA Pol I (Pol Ii), which fully rescues CENP-A localization on replicated *X. borealis* chromosomes. p-values (top to bottom) by one-way ANOVA with Tukey post-hoc analysis: 0.9794, 0.7979, 0.0005. (**D**) Percentage of mitotic chromosomes with ultrathin regions in *X. tropicalis* extracts treated with solvent control or 1 µM BMH-21 (Pol Ii). Pol I inhibition also rescues *X. borealis* chromosome morphology defects. p-values (top to bottom) by one-way ANOVA with Tukey post-hoc analysis: 0.5078, 0.9999, 0.0469. (**E**) Percentage of chromosomes with centromeric CENP-A staining in *X. tropicalis* extracts treated with solvent control or 25 µM triptolide to inhibit RNA Pol II (Pol IIi). *X. laevis* chromosomes are partially rescued, while *X. tropicalis* and *X. borealis* chromosomes are not affected. Quantification with N = 3 extracts, N > 322 chromosomes per extract. p-values (top to bottom, then left to right) by one-way ANOVA with Tukey post-hoc analysis: 0.4785, 0.8797, 0.0052, 0.0125, 0.0003, 0.9999. A, B: DNA in cyan, Pol I in red. Scale bar is 5 µm. C, D: N = 3 extracts, N > 172 chromosomes per extract. C-E: ns, not significant.

To test whether RNA Pol I occupancy at rDNA of *X. borealis* chromosomes contributed to the observed defects, *X. tropicalis* extract reactions were treated with the inhibitor BMH-21, which has been shown to dissociate the polymerase from chromatin (Peltonen et al., 2014; Colis et al., 2014). Strikingly, *X. borealis* chromosome morphology defects as well as CENP-A localization were rescued (Fig. 5C, D). Together, these data suggest that the replication stress experienced by *X. borealis* mitotic chromosomes occurs at rDNA loci, and that defects in rDNA chromatin dynamics act to destabilize a subset of *X. borealis* centromeres. In contrast, centromere formation on *X. laevis* chromosomes was not rescued by RNA Pol I inhibition (Fig. S5A, B), further indicating differences in the mechanisms underlying their incompatibility with *X. tropicalis*. However, we observed that inhibition of RNA polymerase II (Pol II) with triptolide partially rescued CENP-A localization to *X. laevis* chromosomes in *X. tropicalis* extract, whereas *X. borealis* chromosomes were not affected (Fig. 5E), and no species’ chromosomes were rescued by inhibition of RNA Pol III (Fig. S5C, D). Therefore, a common theme in hybrid incompatibility among *Xenopus* species may be replication-transcription conflicts that contribute to eviction of CENP-A from a subset of mitotic chromosomes. However, whereas this occurs at rDNA on *X. borealis* chromosomes and depends on RNA Pol I, *X. laevis* defects are driven, at least in part, by RNA Pol II-induced defects. These observations lead to the model that epigenetic mechanisms promoting CENP-A incorporation at centromeres are disrupted by the presence or activity of RNA polymerases that cause under-replication at specific chromosome loci. Whereas *X. laevis* defects can be overcome by driving CENP-A incorporation at centromeres, *X. borealis* defects can only be rescued by blocking replication stress at rDNA, either by preventing fork collapse or by removing RNA Pol I.

### Chromosome mis-segregation can be reduced in hybrid embryos, but inviability persists

To determine whether the incompatibility mechanisms identified through this work are responsible for hybrid inviability in vivo, we performed rescue experiments on cross-fertilized embryos. In vitro translated, paternally-matched CENP-A and HJURP proteins were microinjected into both blastomeres of the 2-cell hybrid embryo produced by fertilizing *X. tropicalis* eggs with *X. laevis* sperm, while *X. tropicalis*/*X. borealis* hybrid embryos were treated with RNA Pol I inhibitor BMH-21. Fewer micronuclei were observed in both cases, indicating a decrease in mitotic errors in hybrid embryos, although not to the low levels seen in wild-type *X. tropicalis* embryos (Fig. 6A, Fig. S6). Thus, the basis of chromosome defects identified using our in vitro egg extract assays also contribute to chromosome segregation defects in vivo. However, despite this partial rescue, treated hybrids died at the same time and in the same manner as untreated sibling controls (Fig. 6B, C, Movie S1, S2). While it is possible that a complete rescue of chromosome segregation defects in the hybrid embryos is required for viability, we predict that other mechanisms that we have not yet identified also contribute, which can be uniquely addressed using a combination of in vitro and in vivo approaches in *Xenopus*.

**FIGURE 6:**
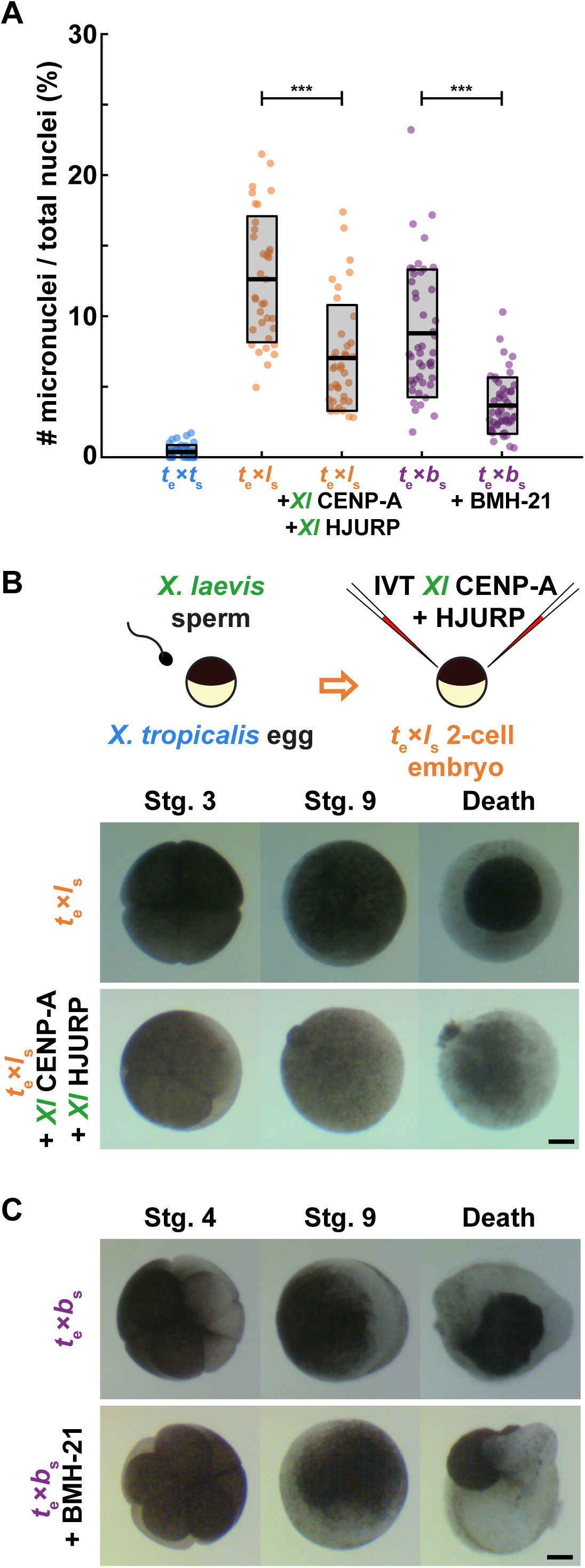
Treatments that rescue CENP-A localization in egg extracts reduce micronuclei formation in hybrid embryos, but inviability persists. (**A**) Quantification of chromosome mis-segregation events as measured by the number of micronuclei compared to total nuclei in treated hybrid embryos. *X. tropicalis* eggs fertilized with *X. laevis* sperm were microinjected with *X. laevis* CENP-A/HJURP, while *X. tropicalis* eggs fertilized with *X. borealis* sperm were treated with Pol 1 inhibitor BMH-21. Embryos were fixed at stage 9 (7 hpf) just before gastrulation and hybrid death. The number of micronuclei was significantly reduced in both cases, but not to control levels measured in *X. tropicalis* eggs fertilized with *X. tropicalis* sperm. N = 3 clutches for each hybrid, N > 15 embryos and > 200 cells per embryo. p-values (left to right) by two-tailed two-sample unequal variance t-tests: 2.111e-7, 2.651e-9; ns, not significant. (**B**) Schematic of experiment and movie frames of *X. tropicalis* eggs fertilized with *X. laevis* sperm microinjected at the two-cell stage with *X. laevis* CENP-A/HJURP. Microinjected hybrid embryos die at the same time and in the same manner as uninjected hybrid controls. N = 10 embryos across 4 clutches. Scale bar is 200 µm. (**C**) Movie frames of *X. tropicalis* eggs fertilized with *X. borealis* sperm that were incubated from the two-cell stage with 1 µM RNA Pol I inhibitor, BMH-21. Treated hybrid embryos die at the same time and in the same manner as untreated hybrid controls. N = 12 embryos across 2 clutches. Scale bar is 200 µm.

## DISCUSSION

Centromeric DNA sequences and centromere and kinetochore proteins have been shown to rapidly co-evolve, which is thought to counteract female meiotic drive and maintain faithful chromosome segregation (Malik and Henikoff, 2001; Malik et al., 2002; Kumon et al., 2021; Pontremoli et al., 2021; Hooff et al., 2017). Our study reveals very low conservation of core centromere DNA sequences across three *Xenopus* species, and differences in protein sequences of *Xenopus* CENP-A and its chaperone HJURP are also observed, indicating co-evolution. However, robust epigenetic mechanisms must operate to maintain centromere compatibility in *Xenopus* hybrids, since many crosses are viable (De Robertis and Black, 1979; Woodland and Ballantine, 1980; Bürki, 1985; Narbonne et al., 2011), and only a subset of chromosomes display centromere/kinetochore defects in inviable hybrids (Gibeaux et al., 2018). Thus, neither differences in centromere sequences nor co-evolved centromere/kinetochore proteins contribute directly to *Xenopus* hybrid inviability.

The *Xenopus* egg extract and sperm chromosome reconstitution system uniquely allowed us to identify mechanisms by which centromere formation is disrupted on specific chromosomes in inviable interspecies hybrids. For *X. tropicalis* eggs fertilized with *X. borealis* sperm, in vitro experiments indicate that defects are due to replication stress at rDNA, since both CENP-A localization and chromosome morphology can be rescued by either preventing replication fork collapse or evicting RNA Pol I (Roussel et al., 1996; Bell et al., 1997; Deng et al., 2019). However, it is unclear why distinct subsets of paternal chromosomes appear to possess ultra-thin regions versus centromere defects. We propose that clustering of repetitive elements including rDNA, pericentromeric, and centromeric repeats during interphase brings together different chromosomal loci and their associated machineries. Normally, such clustering is observed at chromocenters, which may function to stabilize centromeres and promote CENP-A deposition in early G1 of the cell cycle (Brändle et al., 2022; Stellfox et al., 2013). Although discrete chromocenters or other nuclear bodies such as nucleoli have not been observed to form in egg extracts, loci interactions may nevertheless occur during interphase that are disrupted in hybrid reactions and affect four specific *X. borealis* chromosomes due to defects in rDNA replication. While addition of excess CENP-A and its chaperone HJURP can rescue centromere assembly on these chromosomes during interphase, the burst of DNA replication that occurs when metaphase-arrested extract is added (Deng et al., 2019) may simultaneously lead to fork breakage and CENP-A loss. Understanding how formation of fragile sites and centromere loss are related will require a complete *X. borealis* genome assembly that includes rDNA and other repetitive sequences.

Our findings highlight the dynamic interplay between machineries that promote and disrupt centromere assembly. For in vitro reactions reconstituting *X. tropicalis* eggs fertilized with *X. laevis* sperm, the disruption does not involve Pol I or replication stress. Centromere defects appear less severe in this hybrid reaction and can be fully rescued by addition of either species-matched or overexpressed CENP-A/HJURP and partially rescued by Pol II eviction, treatments that may reinforce epigenetic machineries that maintain centromeres. Thus, distinct mechanisms underlie centromere disruption in the two inviable hybrids, but defects in both cases are consistent with observations that aberrant polymerase occupancy or transcription adjacent to the centromere can compromise its assembly (Rošić et al., 2014; Grenfell et al., 2016; Bobkov et al., 2018).

An open question is how the incompatibilities we have characterized in vitro manifest in hybrid embryos in vivo. Whole genome sequencing of the *X. tropicalis* egg/*X. laevis* sperm hybrid just prior to embryo death combined with preliminary Hi-C analysis indicates that the long arms of chromosomes 3L and 4L have been largely eliminated, but the centromere persists on the short arm allowing it to be retained (Gibeaux et al., 2018 and unpublished data). One possible explanation is that under-replication of repetitive sequences adjacent to the centromere in this hybrid initially disrupts centromere assembly, but after chromosome breakage, the adjacent, troublesome sequences are removed and the centromere stabilizes on the short arm while the long arm lacking the centromere frequently ends up in micronuclei and is eventually degraded. Because micronuclei are observed throughout embryogenesis in both inviable hybrids (Gibeaux et al., 2018), multiple rounds of chromosome mis-segregation and instability likely occur that give rise to the terminal karyotype. In the *X. tropicalis* egg/*X. borealis* sperm inviable hybrid that experiences replication stress, a pathway involving p97-mediated extraction and degradation of the replicative helicase that leads to fork breakage and microhomology-mediated end joining events likely operates, which has been well characterized in *Xenopus* egg extracts (Deng et al., 2019). Detailed genomic analysis of chromosome deletions and rearrangements in hybrid embryos will shed light on how replication-transcription conflicts give rise to specific chromosome defects, while additional in vitro experiments will reveal underlying molecular mechanisms.

Death of inviable *Xenopus* hybrids occurs at gastrulation when the zygotic genome undergoes widespread transcriptional activation, and the distinct death phenotypes observed upon fertilization of *X. tropicalis* eggs with either *X. laevis* or *X. borealis* sperm may be due to the different sets of genes affected by the loss of specific chromosomal loci. However, despite a reduction in micronuclei upon hybrid embryo treatments that rescued centromere formation in vitro, death was not delayed or the phenotypes altered in any way. Therefore, we hypothesize that other incompatibilities also contribute to hybrid inviability. In particular, mismatches between mitochondrial and nuclear-encoded genes have been shown to underlie inviability in some hybrids (Lee et al., 2008; Ma et al., 2016).

In conclusion, our findings identify defects in epigenetic centromere maintenance that contribute to hybrid inviability. The combination of in vivo, in vitro, and genomic approaches possible in *Xenopus* promise to provide further mechanistic insights into the molecular basis of hybrid fates and speciation.

## ACKNOWLEDGEMENTS

We thank Daniel Rokshar, Austin Mudd, Sofia Medina-Ruiz, and Mariko Kondo for early access to *X. borealis* CENP-A, CENP-C, HJURP, and H3 sequences. We also thank students Elizabeth Turcotte, Costa Bartolutti, Justin Peng, and Christian Erikson for help with experiments to MK. We are grateful to the Welch, King, Dernberg, Karpen, Lewis, and Rokshar laboratories at UC Berkeley for sharing reagents, discussions, and expertise. We thank all past and present members of the Heald laboratory, Coral Y. Zhou, Gary Karpen, Dirk Hockemeyer, Rasmus Nielsen, and Mark J. Khoury for continuous support and fruitful discussions. M.K. was supported by a National Science Foundation (NSF) GRFP fellowship. O.K.S. was supported by a National Institutes of Health (NIH) T32 GM113854-02 and an NSF GRFP fellowship. A.F.S. was supported by NIH NIGMS R01 GM074728. R.H. was supported by NIH MIRA grant R35 GM118183 and the Flora Lamson Hewlett Chair.

## AUTHOR CONTRIBUTIONS

Conceptualization, Funding acquisition: RH, MK

Methodology, Investigation, Visualization: MK, OKS

Supervision: AFS, RH

Manuscript preparation: MK, RH, OKS, AFS

## SUPPLEMENTARY MATERIALS

Materials and Methods

Figs. S1 to S6

Movies S1 and S2

## MATERIALS AND METHODS

### Lead Contact and Materials Availability

All data and materials are available upon request. Further information and requests for resources and reagents should be directed to Rebecca Heald.

### Experimental Model and Subject Details

All frogs were used and maintained in accordance with standards established by the UC Berkeley Animal Care and Use Committee and approved in our Animal Use Protocol. Mature *Xenopus laevis*, *X. tropicalis*, and *X. borealis* frogs were obtained from Nasco (Fort Atkinson, WI) or the National *Xenopus* Resource (Woods Hole, MA). *Xenopus* frogs were housed in a recirculating tank system with regularly monitored temperature and water quality (pH, conductivity, and nitrate/nitrite levels). *X. laevis* and *X. borealis* were housed at 20-23°C, and *X. tropicalis* were housed at 23-26°C. All animals were fed Nasco frog brittle.

### Chemicals

Unless otherwise states, all chemicals were purchased from Sigma-Aldrich, St. Louis, MO.

### Frog care

*X. laevis*, *X. tropicalis*, and *X. borealis* females were ovulated with no harm to the animals with a 6-, 3-, and 4-month rest interval, respectively, as previously described (Kitaoka et al., 2018). To obtain testes, males were euthanized by over-anesthesia through immersion in ddH_2_O containing 0.15% MS222 (Tricaine) neutralized with 5 mM sodium bicarbonate prior to dissection, and then frozen at -20°C.

### CENP-A ChIP-seq and data analysis

CENP-A MNase ChIP-seq was performed as previously described (Smith et al., 2021). Briefly, livers were extracted from adult *X. borealis* animals and flash frozen. Upon thawing, livers were diced on ice, rinsed in PBS, and buffer 1 (2.5 mM EDTA, 0.5 M EGTA, 15 mM Tris-HCl pH 7.4, 15 mM NaCl, 60 mM KCl, 15 mM sodium citrate 0.5 mM spermidine, 0.15 mM spermine, 340 mM sucrose, supplemented with 0.1 mM PMSF) was added and the tissue dounced using pestle A 12 times. A syringe with 18-gauge needle was backfilled with nuclei mixture and expelled into 2 mL tubes with additional buffer 1. Nuclei were spun at 6,000g for 5 min at 4°C, and washed 3 times with buffer 3 (2.5 mM EDTA, 0.5 M EGTA, 15 mM Tris-HCl pH 7.4, 15 mM NaCl, 60mM KCl, 15 mM sodium citrate 0.5 mM spermidine, 0.15 mM spermine, 340 mM sucrose, supplemented with 0.1 mM PMSF). Nuclei quality was checked and nuclei were counted by hemocytometer. ∼5-10 million nuclei were used per IP reaction.

For MNase digestion, CaCl_2_ was added to each reaction tube to 5 mM together wtih 300 U of MNase. Digestion was performed at RT for 30 min and reaction was quenched with 10 mM EDTA and 5 mM EGTA. Nucei were lysed with 0.05% IGEPAL CA-630 in ice for 10 min. Following an initial spin 1,500g 5 min 4°C, the pellet was resuspended in 500 µL buffer 3 + 200 mM NaCl and rotated overnight at 4°C to extract mononucleosomes. Samples were precleared, input fractions were taken and CENP-A mononucleosomes were isolated with 10 µg rabbit anti *X. laevis* CENP-A antibody prebound to protein A dynabeads in 200 µL TBST with rotation overnight at 4°C. Beads were washed and eluted, mononucleosomal DNA was isolated with Ampure beads, and sequencing libraries were prepared using NEBNext fit for Illumina sequencing which was performed on a NovaSeq instrument with paired end 150bp sequencing.

*X. laevis* and *X. tropicalis* CENP-A CHIP-seq datasets were used from previously described studies (Smith et al., 2021; Bredeson et al., 2021). CENP-A ChIP and Input libraries from each species were processed to identify CENP-A enriched *k*-mers using the *k*-mer counting pipeline that normalizes *k*-mer counts by sequencing depth of each library (https://github.com/straightlab/xenla-cen-dna-paper). For this study 25bp *k*-mers were used and kmc was run with ci=10, indicating that *k*-mers must be found 10 times in the dataset to be considered. This was chosen so that more *k*-mers were identified from each species to make comparisons more likely.

A phylogram was generated using a method similar to that previously described (Smith et al., 2021). From each species full length ChIP-seq reads were selected based on the presence of at least 80 CENP-A enriched *k*-mers. The reads from each species that met this criterion were then clustered by sequence similarity using cd-hit-est (Fu et al., 2012) using sequential rounds of clustering by 98%, 95%, and 90% identical by sequence. The 20 top clusters from each species were then selected for phylogram generation using Geneious (7.1.4) Tree Builder with the following settings: Genetic Distance Model=Tamura-Nei, Tree building method=Neighbor-joining, Outgroup=No outgroup, Alignment Type=Global alignment, Cost Matrix=93% similarity. Colors for each species were added manually.

### Protein sequence alignments

Multiple sequence alignments were performed using Clustal Omega (default parameters). Sequence similarities were determined by pair-wise alignments using EMBOSS Needle (default parameters).

### *Xenopus* egg extracts

*X. tropicalis* metaphase-arrested egg extracts and spindle reactions were prepared as previously described (Hannak and Heald, 2006; Brown et al., 2007; Maresca and Heald, 2006). Briefly, freshly laid, metaphase II-arrested eggs were collected, dejellied, packed and crushed by centrifugation. The cytoplasmic layer was collected with a syringe and 18G needle, then supplemented with 10 µg/mL of leupeptin, pepstatin, and chymostatin (LPC), 20 µM of cytochalasin B, and energy mix (3.75 µM creatine phosphate, 0.5 µM ATP, 0.5 µM MgCl_2_, 0.05 µM EGTA). Typical reactions contained 20 µL CSF extract, sperm nuclei at a final concentration of 500 nuclei/µL, and rhodamine-labeled porcine brain tubulin at a final concentration of 50 µg/mL.

### Chromosome immunofluorescence

Spindle reactions were prepared, spun-down, and processed for immunofluorescence as previously described (Hannak and Heald, 2006; Maresca and Heald, 2006). Briefly, the extract reactions were fixed for 5-10 min with 2% formaldehyde and spun down at 5,500 rpm (5821.9 x g) for 20 min at 16°C. The coverslips were incubated for 30 s in cold methanol, washed in PBS + 0.1% NP40, and blocked overnight in PBS + 3% BSA at 4°C. We used rabbit anti-xCENPA, 1:500 (Milks et al., 2009; Moree et al., 2011), mouse anti-myc (9E10 clone, 1:500), rabbit anti-POLR1A (Novus Biologicals, 1:500), and mouse anti-UBTF (Abnova, 1:500) antibodies. Primary antibodies were added for 1 h in PBS + 3% BSA. After washing with PBS + 0.1% NP40, the coverslips were incubated with 1:1000 anti-rabbit or mouse secondary antibodies coupled to Alexa Fluor 488 or 568 (Invitrogen), respectively, for 30 min and then with 1:1000 Hoechst (Invitrogen) for 5 min. The coverslips were then washed and mounted for imaging with Vectashield (Vector Labs). Each presented dataset was obtained from three independent egg extracts.

### Nuclear DNA FISH for FCR centromeric sequences

Nuclear DNA FISH using probes against various FCR monomers was performed as previously described (Smith et al., 2021). Briefly, pJET1.2 plasmids containing 150 bp FCR monomer sequences were PCR-amplified and fluorescently labeled using random hexamer priming and Klenow (exo-) polymerase (New England Biolabs). Both Alexa Fluor 488 and 568 dUTP-conjugated fluorophores were used. Probes were desalted to remove unincorporated nucleotides, then precipitated and cleaned before resuspension in hybridization buffer (65% formamide, 5X SSC, 5X Denhardts with 1% blocking reagent (Roche), 0.5 mg/mL salmon sperm DNA added fresh). Each experiment used 4 uL of probe mixed with 4 uL of hybridization buffer.

Nuclei were assembled in egg extract, spun down onto coverslips, and probed with CENP-A antibody as previously described in (Levy and Heald, 2010) and detailed above. Samples proceeded to FISH by fixation in 2.5% formaldehyde in PBS for 10 min, washed in PBS, and dehydrated with increasing concentrations of 70-100% ice-cold ethanol. Coverslips were blocked for 30 m in hybridization buffer. Probes were warmed and mixed with hybridization buffer before being added to samples, flipping coverslips onto glass slides for hybridization. These “sandwiches” were incubated at 80°C for 10 min, then incubated overnight at 37°C. Coverslips were removed from glass slides carefully with 4X SSC, washed thoroughly in SSC, stained with Hoechst and mounted with Vectashield (Vector Labs).

### Protein expression in reticulocyte lysate

To generate plasmids for expression of species-specific *X. laevis*, *X. tropicalis*, and *X. borealis* CENP-A, total RNA was isolated from stage 9 embryos. Embryos were homogenized mechanically in TRIzol (Thermo Fisher Scientific) using up to a 30-gauge needle and processed according to manufacturer’s instructions. After resuspension in nuclease-free H2O, RNAs were cleaned using a RNeasy kit (Qiagen) according to manufacturer’s instructions, and cDNA was synthesized using the SuperScript III First Strand Synthesis system (Thermo Fisher Scientific) according to the manufacturer’s instructions. The *X. laevis*, *X. tropicalis*, and *X. borealis* CENP-A sequences were then PCR-amplified from the cDNA. The amplified sequence was then subcloned into a pCS2+ vector using Gibson assembly. The constructs were then amplified using XL1-Blue competent *E. coli* (Agilent).

The TnT Sp6-coupled rabbit reticulocyte system (Promega) was used for in vitro transcription/translation (IVT) of plasmid DNA according to the manufacturer’s protocol. 2-10% of the final egg extract reaction volume was added prior to addition of sperm nuclei; for CENP-A, this corresponds to 8-80 times endogenous protein levels.

### Western blots

Increasing volumes of egg extracts and reticulocyte lysate were subject to SDS-PAGE and wet transferred to PVDF membranes. Blots were blocked with PBS + 0.1% Tween + 5% milk for 1 h, probed with primary antibodies diluted in PBS + 0.1% Tween + 5% milk for 1 h, rinsed 3x over a 10 m period with PBS + 0.1% Tween, then probed with secondary antibodies (Rockland Immunochemicals; goat anti-rabbit DyLight 800 and donkey anti-mouse DyLight 680, 1:10,000) diluted in PBS + 0.1% Tween for 30 m. Blots were scanned on an Odyssey Infrared Imaging System (Li-Cor Biosciences). Band intensities were quantified using FIJI.

### Drug treatments

*X. tropicalis* extract was supplemented with the following drugs and concentrations: Aphidicolin (DNA replication inhibitor, 10 µg/mL), BMH-21 (RNA Polymerase I inhibitor, 1 µM), NMS-873 (p97 inhibitor, 10 µM), MLN-8237 (Aurora A inhibitor, 1 µM, Selleck Chemicals), BI-2536 (Polo kinase 1 inhibitor, 1 µM, Selleck Chemicals), Triptolide (RNA Polymerase II inhibitor, 25 µM)

### Chromosome and nuclei imaging

Chromosomes were imaged using Micromanager 1.4 software (Edelstein et al., 2014) and nuclei were imaged using Olympus cellSens Dimension 2 software on an upright Olympus BX51 microscope equipped with an ORCA-ER or ORCA-Spark camera (Hamamatsu Photonics) and Olympus UPlan 60X/NA 1.42 oil objective. All images across all datasets were taken using the same exposure settings.

### In vitro fertilization and cross-fertilizations

In vitro fertilization and cross-fertilizations were performed as previously described (Gibeaux et al., 2018; Gibeaux and Heald, 2019; Kitaoka et al., 2018). *X. laevis*, *X. borealis*, and *X. tropicalis* males were injected with 500, 300, and 250 U, respectively, of human chorionic gonadotropin hormone (hCG) 12-24 h before dissection. Testes were collected in Leibovitz L-15 Medium (Gibco, Thermo Fisher Scientific) supplemented with 10% fetal bovine serum (FBS; Gibco) for immediate use. *X. tropicalis* females were primed with 10 U of hCG 12-18 h before use and boosted with 250 U of hCG on the day of the experiment. As soon as the first eggs were laid (∼3 h after boosting), the male was euthanized and dissected. Two *X. tropicalis* testes or one *X. laevis* or *X. borealis* testis were added to 1 mL of L-15 + 10% FBS. *X. tropicalis* females were squeezed gently to deposit eggs onto glass Petri dishes (Corning) coated with 1.5% agarose in 1/10X MMR (1X MMR: 100 mM NaCl, 2 mM KCl, 2 mM CaCl2, 1 mM MgSO4, 0.1 mM EDTA, 5 mM HEPES-NaOH pH 7.6) Testes were homogenized using a pestle in L-15 + 10% FBS to create sperm solution. Any liquid in the Petri dishes was removed, and the eggs were fertilized with 500 uL of sperm solution per dish. Eggs were swirled in the solution to separate them and incubated for 5 min with the dish slanted. Dishes were flooded with ddH_2_O and incubated for 10 min. ddH_2_O was exchanged for 1/10X MMR and incubated for 10 min. The jelly coats were removed with a 3% cysteine solution (in ddH_2_O-NaOH, pH 7.8). After extensive washing with 1/10X MMR (at least four times), embryos were incubated at 23°C until the first cleavage at 1 hour post fertilization (hpf). Fertilized embryos were then sorted and placed in a mesh-bottomed dish for microinjection as described below.

### Embryo microinjection

At stage 2 (2-cell embryo), embryos were transferred to a 1/9X MMR + 3% Ficoll. IVT reticulocyte lysate was backloaded into a needle pulled from a 1 mm glass capillary tube (TW 100F-4, World Precision Instruments) using a P-87 Micropipette Puller (Sutter Instrument). Embryos were placed in a mesh-bottomed dish and microinjected in both blastomeres with 2 nL of the IVT reticulocyte lysate using a Picospritzer III microinjection system (Parker) equipped with a MM-3 micromanipulator (Narishige). Injected embryos were transferred to a new dish and incubated at 23°C in 1/9X + 3% Ficoll for several hours, then buffer exchanged for 1/10X MMR overnight.

### Embryo video imaging

Imaging dishes were prepared using an in-house PDMS mold designed to print a pattern of 0.9 mm large wells in agarose that allowed us to image six *X. tropicalis* embryos simultaneously within the 3 mm x 4 mm camera field of view for each condition. Embryos were imaged from stage 3 after microinjection. Treatment and control videos were taken simultaneously using two AmScope MD200 USB cameras (AmScope), each mounted on an AmScope stereoscope. Time-lapse movies were acquired at a frequency of one frame every 10 s for 20 h and saved as Motion JPEG using a Matlab (The MathWorks) script. Movie post-processing (cropping, concatenation, resizing, and addition of scale bar) was done using Matlab and FIJI (Schindelin et al., 2012). All Matlab scripts written for this study are available upon request. Two of the scripts used here were obtained through the MATLAB Central File Exchange: ‘videoMultiCrop’ and ‘concatVideo2D’ by ‘Nikolay S’.

### Embryo whole-mount immunofluorescence

Embryos were fixed at the desired stages for 1-3 h using MAD fixative (2:2:1 methanol: acetone: DMSO). After fixation, embryos were dehydrated in methanol and stored at - 20°C. Embryos were then processed for immunofluorescence as previously described (Gibeaux et al., 2018). Briefly, embryos were gradually rehydrated in 0.5X SSC (1X SSC: 150 mM NaCl, 15 mM Na citrate, pH 7.0), then bleached with 2% H_2_O_2_ in 0.5X SSC with 5% formamide for 2 h under light. Embryos were washed with PBT (1X PBS, 0.1% Triton X-100, 2 mg/mL bovine serum albumin). Embryos were blocked in PBT supplemented with 10% goat serum and 5% DMSO for 1-3 h and incubated overnight at 4°C in PBT supplemented with 10% goat serum and primary antibodies. We used mouse anti-beta-tubulin (E7, Developmental Studies Hybridoma Bank, 1:300 dilution) and rabbit anti-histone H3 (Abcam, 1:500 dilution). Embryos were then washed 4 x 2 h in PBT and incubated overnight at 4°C in PBT supplemented with goat anti-mouse and goat anti-rabbit secondary antibodies coupled to Alexa Fluor 488 and 568 (Invitrogen). Embryos were then washed 4 x 2 h in PBT and gradually dehydrated in methanol. Finally, embryos were cleared in Murray’s clearing medium (2:1 benzyl benzoate: benzyl alcohol). Embryos were placed in a reusable chamber (Thermo Fisher) for confocal microscopy.

### Confocal microscopy

Confocal microscopy was performed on an inverted Zeiss LSM 800 using the Zeiss Zen software, a Plan-Achromat 20X/0.8 air objective and laser power 0.5-2%, on multiple 1024×1024 pixel plans spaced of 0.68 µm in Z. Images are mean averages of two scans with a depth of 16 bits. Pinhole size always corresponded to 1 Airy unit.

### Quantification and Statistical Analysis

Quantification of CENP-A localization on mitotic chromosomes was determined manually in a dataset of 100 images from one extract. Quantification of ultra-thin chromosomal regions was also determined manually in parallel from the same datasets. Only single chromosomes were counted. Each dataset had ∼150-400 chromosomes. The average of each extract was calculated as a percentage of total chromosome number. Averages were plotted in Matlab, and statistical significance and p-values were determined with two-tailed, two-sample unequal variance t-tests or one-way ANOVA with Tukey post-hoc analysis in Microsoft Excel. The number of egg extracts used, individual chromosomes counted, and p-values are listed in the figure legends. For all box plots, the thick line inside the box indicates the average across biological replicates, and the upper and lower box boundaries indicate the standard deviation.

PolR1A and UBF fluorescent intensity on *X. borealis* ultra-thin chromosomes were quantified in FIJI by measuring the intensity of the stretched region specifically and comparing it to a random non-stretched region on the same chromosome. All intensity measurements were normalized to the samples’ Hoechst intensity.

Micronuclei in embryos were quantified at the relevant stages as the number of observed micronuclei divided by the number of nuclei, counted manually in FIJI. Statistical significance was determined by two-tailed, two-sample unequal variance t-tests.

---

**Figure S1:**
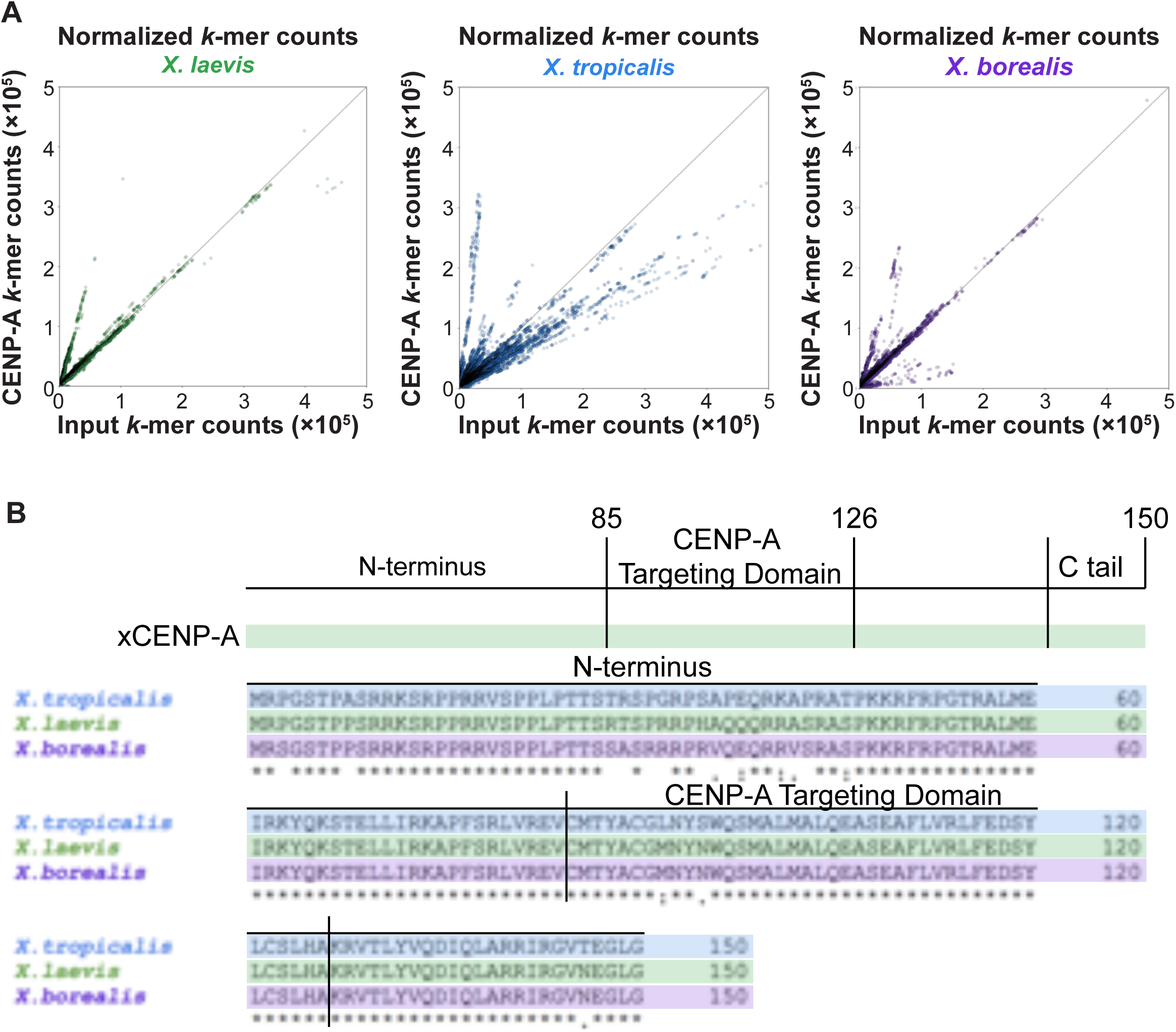
Comparison of centromere DNA and CENP-A protein sequences. (**A**) Scatter plots of normalized *k*-mer counts from Input and CENP-A ChIP-seq sequencing libraries from *X. laevis*, *X. tropicalis*, and *X. borealis*. The dotted line (x=y) indicates *k*-mers that are equally abundant in both libraries. *k*-mer counts reveal distinct patterns in the three species. (**B**) Protein sequence alignment comparing CENP-A across the three *Xenopus* species. Differences are observed in the N-terminal region and the CENP-A targeting domain (CATD).

**FIGURE S2:**
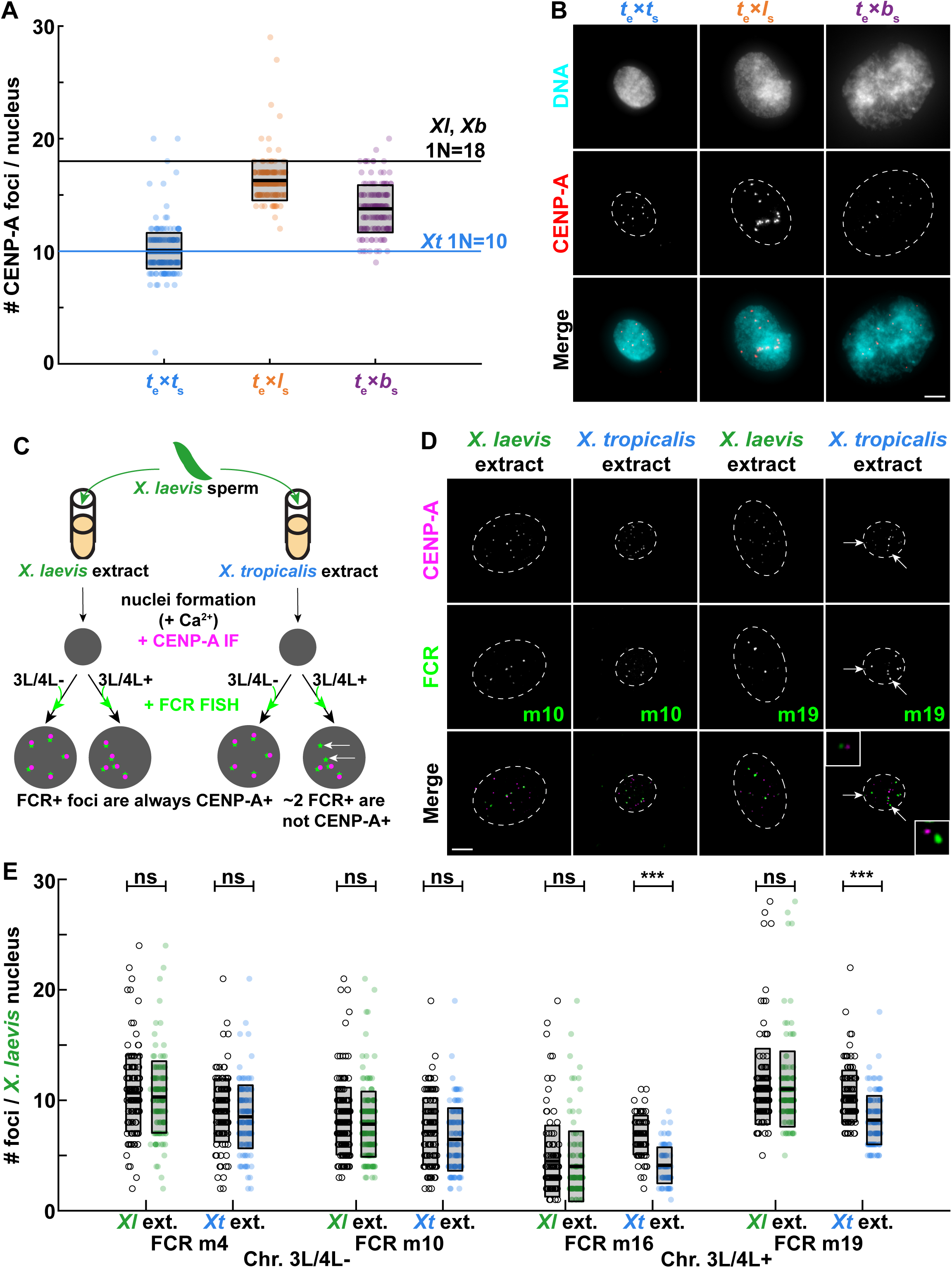
CENP-A is lost from *X. laevis* chromosomes 3L and 4L. (**A**) Quantification of the number of CENP-A foci in interphase nuclei assembled in *X. tropicalis* egg extract. Whereas *X. tropicalis* nuclei on average possess 10 foci corresponding to the 10 sperm chromosomes, *X. laevis* and *X. borealis* interphase nuclei possess an average of 16 and 14 CENP-A foci, respectively, which does not match the 18 sperm chromosomes of these species. Quantification with N = 3 extracts, N > 64 nuclei per extract. (**B**) Representative images of *X. tropicalis*, *X. laevis* and *X. borealis* nuclei formed in *X. tropicalis* extract. DNA in cyan, CENP-A in red. Scale bar is 5 µm. (**C**) Experimental schematic for specific centromere quantification. *X. laevis* sperm nuclei were cycled into interphase in either *X. laevis* or *X. tropicalis* egg extract. All centromeres were detected by CENP-A immunofluorescence and a subset of centromeres were identified by FCR (frog centromeric repeat) FISH (Smith et al., 2021). Probes prepared from two sequences not present in centromeres of chromosomes 3L or 4L (3L/4L-= m4, m10), were compared with probes made using two sequences present in centromeres of chromosomes 3L and 4L (3L/4L+ = m16, m19). In *X. laevis* extract, all FCR+ foci should co-localize with CENP-A, as 18/18 centromeres are maintained. If centromeric CENP-A staining is lost specifically from chromosomes 3L and 4L in *X. tropicalis* extract, 2 3L/4L+ FCR+ foci should not colocalize with CENP-A (panel A). (**D**) Images of *X. laevis* sperm nuclei formed in *X. laevis* or *X. tropicalis* extract probed by FISH for FCR monomer m10 or m19 (green) and by immunofluorescence of CENP-A (magenta). Insets show the 2 m19 FCR+ foci not co-localized with CENP-A, while all other FCR+ foci co-localize with CENP-A. DNA periphery is marked by the dashed white lines. Scale bar is 5 µm. (**E**) Quantification of CENP-A foci that co-localize with FCR+ foci in *X. laevis* vs. *X. tropicalis* extract. In *X. laevis* extract, all FCR+ foci co-localize with CENP-A. However, in *X. tropicalis* extract, ∼2 m16 or m19 FCR+ foci do not co-localize with CENP-A, corresponding to the loss of CENP-A localization on chromosomes 3L and 4L. Quantification with N = 2 extracts, N > 50 nuclei and > 800 centromeres per probe per extract. p-values by two-tailed two-sample unequal variance t-tests (left to right): 0. 3562, 0.0916, 0.3708, 0.0499, 0.2426, 2.797e-19, 0.5485, 7.972e-13; ns, not significant. Open circles are FCR+ foci, filled circles represent foci that are both FCR+ and CENP-A+.

**Figure S3:**
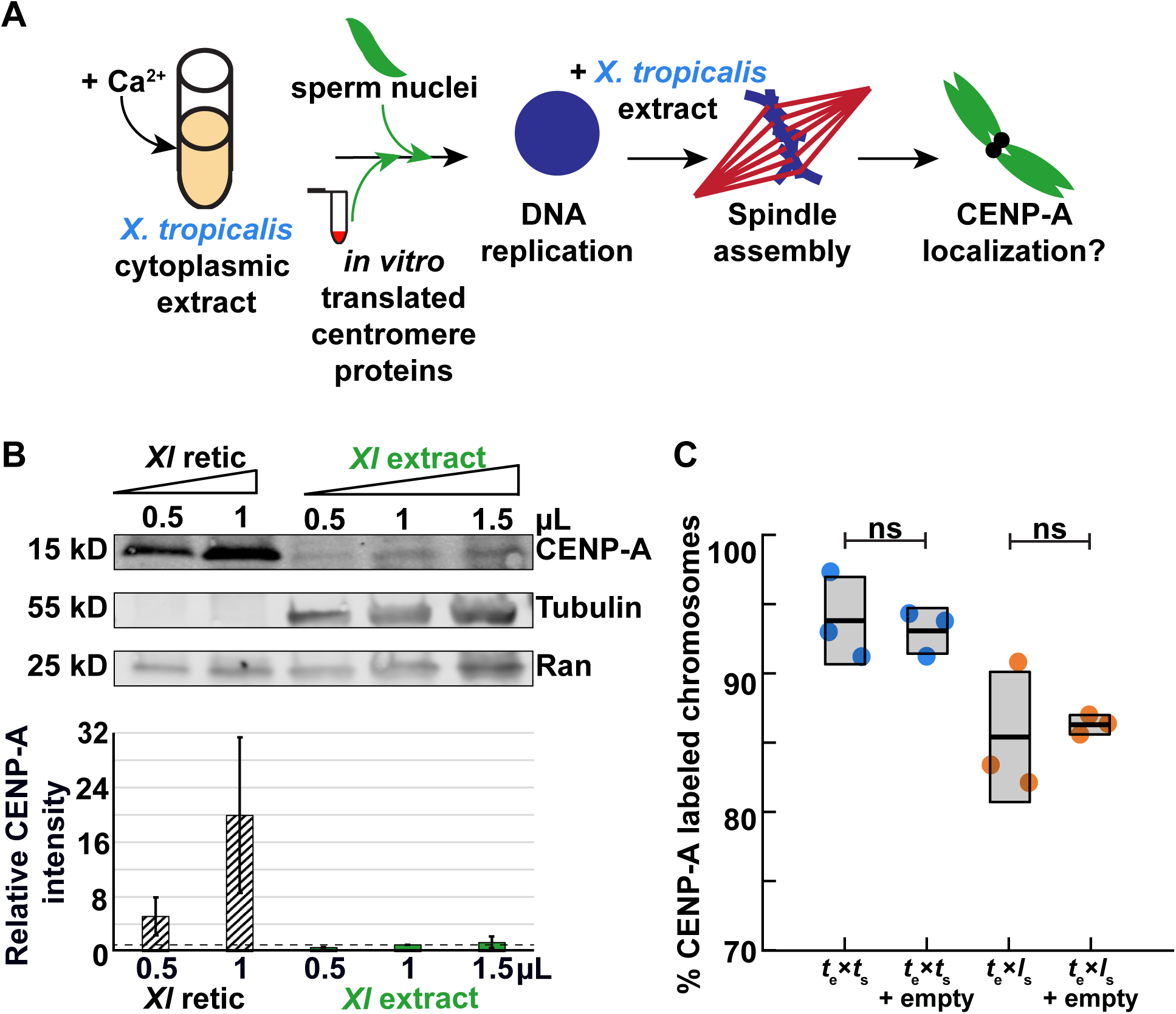
Driving CENP-A assembly with proteins expressed in reticulocyte lysate. (**A**) Experimental schematic of extract reactions in which reticulocyte lysate is added at the onset of interphase to mimic the timing of CENP-A deposition in G1. (**B**) Representative Western blot of *X. laevis* CENP-A protein expressed in reticulocyte lysate and quantification of three blots showing band intensity normalized to CENP-A levels in 1 µL of *X. laevis* egg extract (dotted line). CENP-A is approximately twenty times more concentrated in reticulocyte lysate compared to *X. laevis* extract, with amounts added to chromosome/nuclear assembly reactions corresponding to between 8 and 80 times endogenous CENP-A levels. (**C**) Percentage of replicated *X. laevis* or *X. tropicalis* chromosomes with CENP-A staining in *X. tropicalis* extract supplemented with unprogrammed reticulocyte lysate. Lysates containing empty expression vectors have no effect on centromere staining. Quantification from N = 3 extracts, N > 298 chromosomes per extract. p-values (left to right) by two-tailed two-sample unequal variance t-tests: 0.7433, 0.7755; ns, not significant.

**FIGURE S4:**
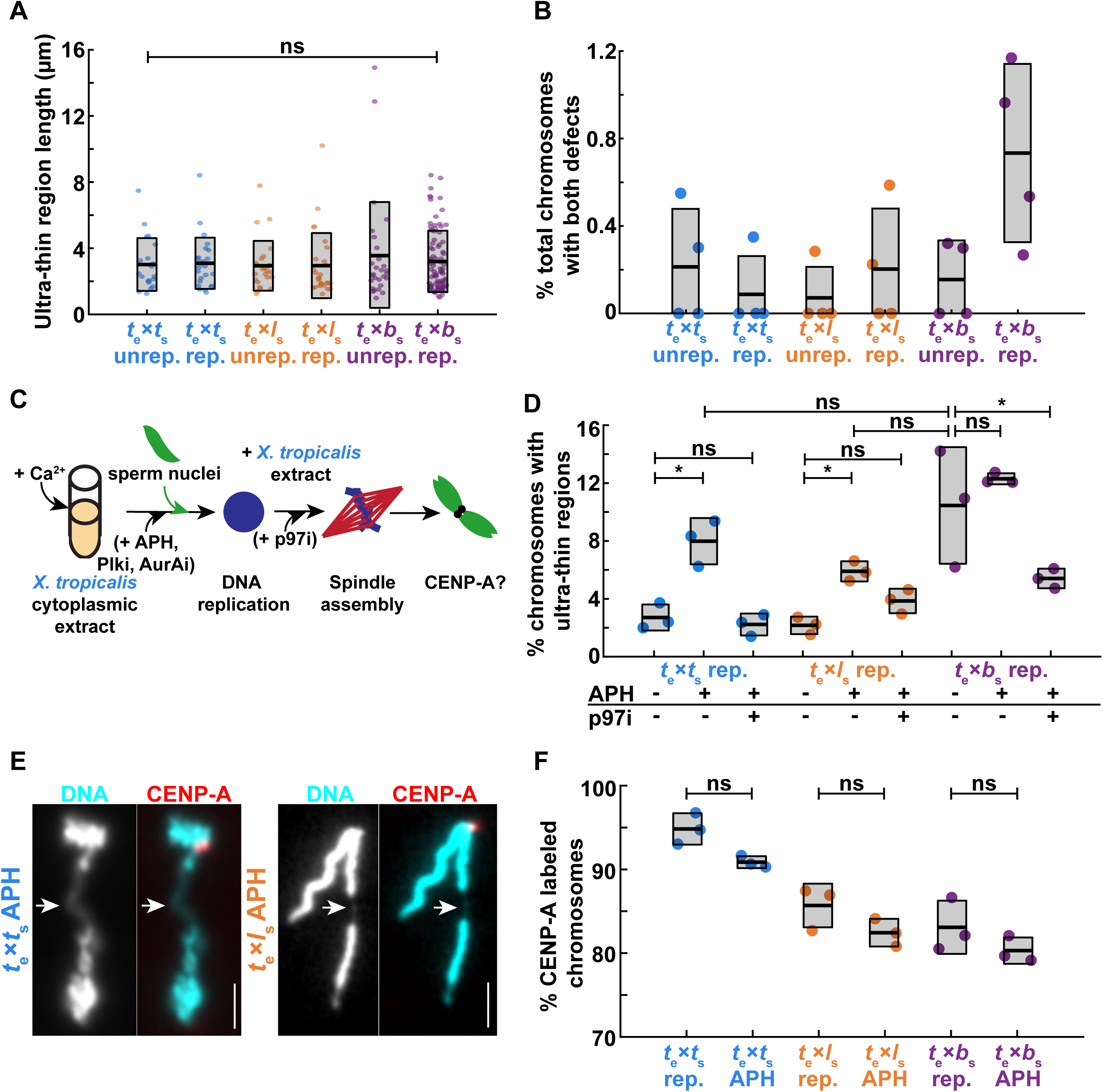
Characterization of chromosome morphology defects that can be induced by aphidicolin and rescued by p97 inhibition. (**A**) Quantification of ultra-thin region lengths, which average ∼2-3 µm on mitotic chromosomes of all three *Xenopus* species. p-value by one-way ANOVA = 0.8712. (**B**) Percentage of *X. tropicalis*, *X. laevis*, and *X. borealis* chromosomes with ultra-thin regions that have also lost CENP-A staining. Only a small fraction of chromosomes with ultra-thin regions also show centromere loss. Across all species, only ∼0.2-0.6% of all chromosomes exhibit both morphological and centromere defects, corresponding to 1-4 chromosomes out of ∼350 total chromosomes per extract. (**C**) Experimental schematic illustrating when inhibitors are added to *X. tropicalis* extract reactions. (**D**) Percentage of mitotic chromosomes with ultrathin regions in *X. tropicalis* extracts treated with solvent control or 10 µg/mL aphidicolin (APH) to inhibit DNA replication, and with or without 10 µM p97 ATPase inhibitor NMS-873 (p97i) to prevent removal of stalled replication forks. Aphidicolin increased the prevalence of ultra-thin chromosome regions on *X. tropicalis* and *X. laevis* chromosomes, but did not significantly exacerbate these regions on *X. borealis* chromosomes. Inhibition of p97 rescued the chromosome morphology defects. Quantification from N = 3 extracts, N > 138 chromosomes per extract. p-values (top to bottom, then left to right) by two-tailed two-sample unequal variance t-tests: 0.6106, 0.0217, 0.9986, 0.8708, 0.9999, 0.9159, 0.0151, 0.0023. (**E**) Representative images of *X. tropicalis* and *X. laevis* mitotic chromosomes following aphidicolin treatment. DNA in cyan, CENP-A in red. Scale bar is 5 µm. (**F**) Percentage of replicated chromosomes with centromeric CENP-A staining in *X. tropicalis* extracts treated with solvent control or 10 µg/mL aphidicolin. Inhibition of DNA replication does not affect centromere formation on any species’ chromosomes. p-values (left to right) by two-tailed two-sample unequal variance t-tests: 0.0523, 0.1554, 0.2679. A, B: N = 3 extracts, N > 20 chromosomes per condition. D, F: N = 3 extracts, N > 150 chromosomes per extract. A, D, F: ns, not significant.

**Figure S5:**
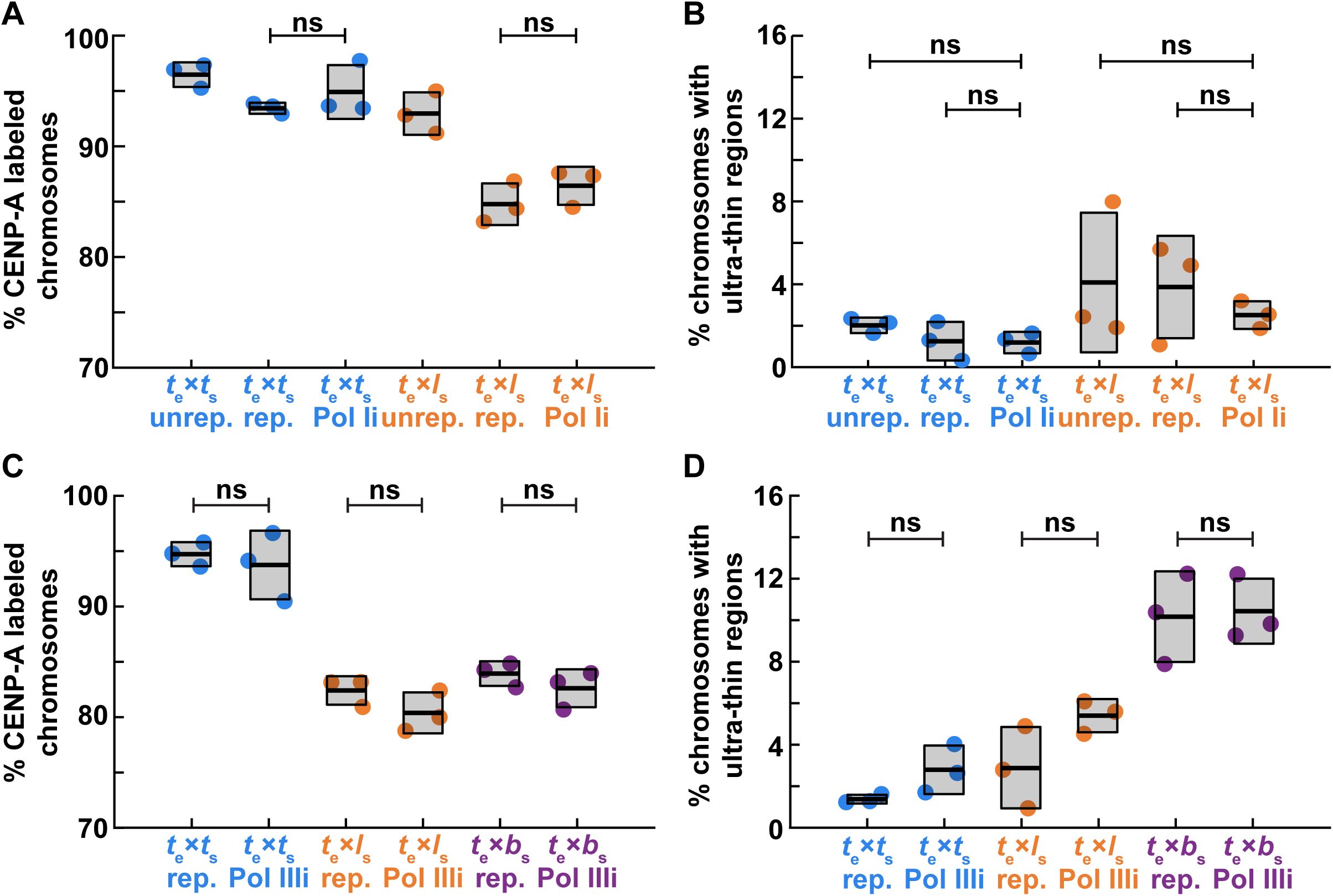
Pol I transcription inhibition does not affect *X. tropicalis* or *X. laevis* chromosomes, while Pol III inhibition had no effect on any species. (**A**, **B**) The percentage of *X. tropicalis* or *X. laevis* mitotic chromosomes formed in *X. tropicalis* egg extract with centromeric CENP-A staining (A) or ultrathin regions (B) is unchanged upon treatment with 1 µM BMH-21 to inhibit RNA Pol I (Pol Ii). Quantification with N = 3 extracts, N > 113 chromosomes per extract. p-values by one-way ANOVA with Tukey post-hoc analysis: (A, left to right) 0.9702, 0.9413, (B, left to right, then top to bottom) 0.9995, 0.9711, 1, 0.9882. (**C**, **D**) The percentage of *X. tropicalis*, *X. laevis*, or *X. borealis* mitotic chromosomes formed in *X. tropicalis* egg extract with centromeric CENP-A staining (C) or ultrathin regions (D) is unchanged upon treatment with 20 µM ML-69218 to inhibit RNA Pol III (Pol IIIi) Quantification with N = 3 extracts, N > 179 chromosomes per extract. p-values by one-way ANOVA with Tukey post-hoc analysis: (C, left to right) 0.9389, 0.7506, 0.9416, (D, left to right) 0.8431, 0.3540, 0.9999. A-D: ns, not significant.

**FIGURE S6:**
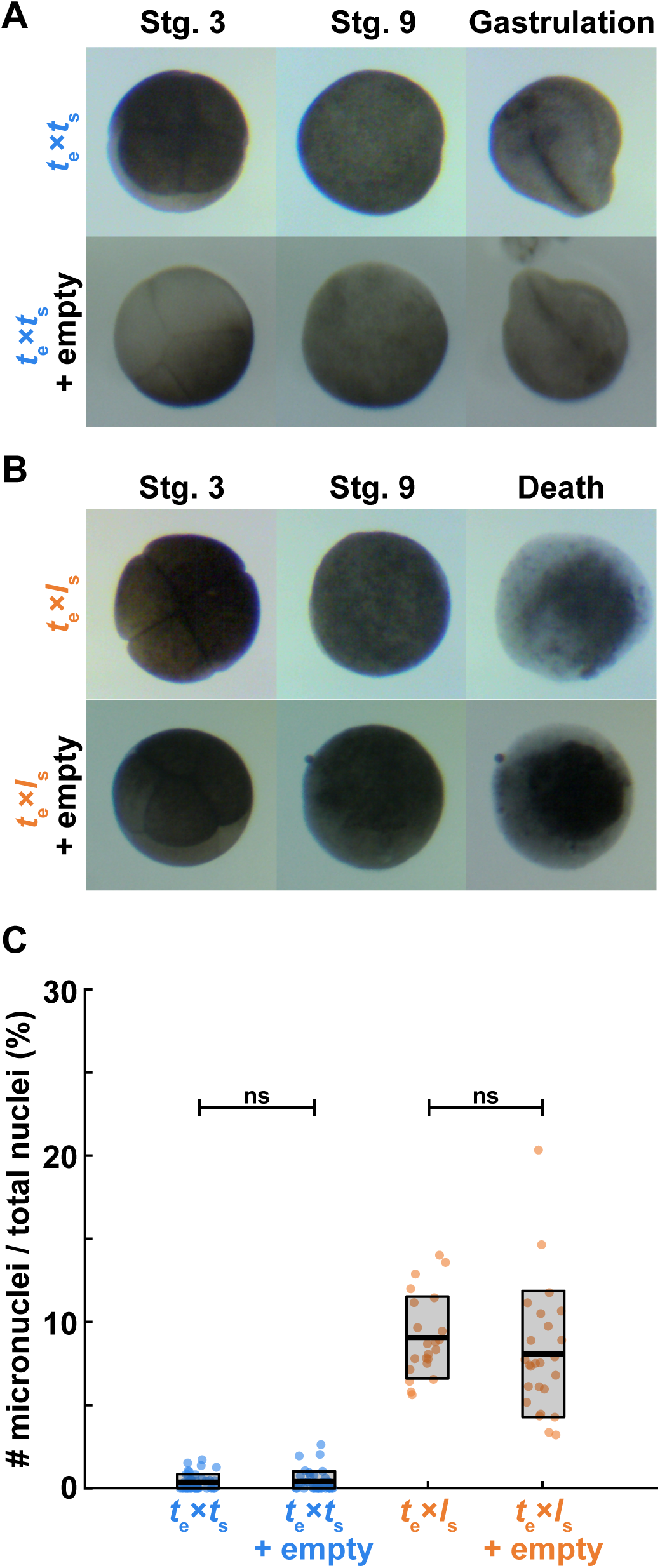
Microinjection with reticulocyte lysate does not affect embryo development or chromosome segregation. (**A**) Movie frames of untreated *X. tropicalis* embryos and embryos microinjected at the two-cell stage with empty reticulocyte lysate into both blastomeres show that embryonic development is not affected by the procedure. N = 15 embryos across 3 clutches. Scale is 200 µm. (**B**) Movie frames of untreated *X. tropicalis/X. laevis* hybrid embryos and hybrid embryos microinjected at the two-cell stage with empty reticulocyte lysate into both blastomeres show that the embryonic death phenotype is not affected by the procedure. N = 12 embryos across 2 clutches. Scale is 200 µm. (**C**) Quantification of the number of micronuclei compared to total nuclei in stage 9 *X. tropicalis* embryos or *X. tropicalis*/*X. laevis* hybrid embryos microinjected with empty reticulocyte lysate. The prevalence of micronuclei is unaffected by the procedure. p-values (left to right) by two-tailed two-sample unequal variance t-tests: 0.749, 0.288; ns, not significant.

**MOVIE S1: Microinjection of *X. laevis* CENP-A and HJURP *in vitro* translated proteins into *t*_e_×*l*_s_ hybrids does not rescue viability.**

*X. tropicalis* eggs were fertilized with *X. laevis* sperm and microinjected with *in vitro* translated paternally-matched CENP-A and HJURP (right) at stage 2. Embryos were imaged in separate dishes from stage 3 in 1/10X MMR. The movie plays 20 h in 15s (rate of 120 frames per second). Scale bar corresponds to 200 µm.

**MOVIE S2: Treatment with RNA Polymerase I inhibitor BMH-21 does not rescue *t*_e_×*b*_s_ hybrid viability.**

*X. tropicalis* eggs were fertilized with *X. borealis* sperm and incubated with 1 µM BMH-21 in 1/10X MMR (right) from stage 2. The movie plays 20 h in 15s (rate of 120 frames per second). Scale bar corresponds to 200 µm.

## REFERENCES

1. Anselm, E., A.W. Thomae, A.A. Jeyaprakash, and P. Heun. 2018. Oligomerization of Drosophila Nucleoplasmin-Like Protein is required for its centromere localization. Nucleic Acids Res. 46:11274–11286. doi:10.1093/nar/gky988.

2. Bell, P., C. Mais, B. McStay, and U. Scheer. 1997. Association of the nucleolar transcription factor UBF with the transcriptionally inactive rRNA genes of pronuclei and early Xenopus embryos. J. Cell Sci. 110:2053–2063.

3. Bell, P., and U. Scheer. 1997. Prenucleolar bodies contain coilin and are assembled in Xenopus egg extract depleted of specific nucleolar proteins and U3 RNA. J. Cell Sci. 110:43–54. doi:10.1242/jcs.110.1.43.

4. Bernad, R., P. Sánchez, T. Rivera, M. Rodríguez-Corsino, E. Boyarchuk, I. Vassias, D. Ray-Gallet, A. Arnaoutov, M. Dasso, G. Almouzni, and A. Losada. 2011. Xenopus HJURP and condensin II are required for CENP-A assembly. J. Cell Biol. 192:569–582. doi:10.1083/jcb.201005136.

5. Blum, J.A., S. Bonaccorsi, M. Marzullo, V. Palumbo, Y.M. Yamashita, D.A. Barbash, and M. Gatti. 2017. The Hybrid Incompatibility Genes Lhr and Hmr Are Required for Sister Chromatid Detachment During Anaphase but Not for Centromere Function. Genetics. 207:1457–1472. doi:10.1534/genetics.117.300390/-/DC1.1.

6. Bobkov, G.O.M., N. Gilbert, and P. Heun. 2018. Centromere transcription allows CENP-A to transit from chromatin association to stable incorporation. J. Cell Biol. 217:1957–1972. doi:10.1083/jcb.201611087.

7. Brändle, F., B. Frühbauer, and M. Jagannathan. 2022. Principles and functions of pericentromeric satellite DNA clustering into chromocenters. Semin. Cell Dev. Biol. doi:10.1016/j.semcdb.2022.02.005.

8. Bredeson, J. V., A.B. Mudd, S. Medina-Ruiz, T. Mitros, O.K. Smith, K.E. Miller, J.B. Lyons, S.S. Batra, J. Park, K.C. Berkoff, C. Plott, J. Grimwood, J. Schmutz, G. Aguirre-Figueroa, M.K. Khokha, M. Lane, I. Philipp, M. Laslo, J. Hanken, G. Kerdivel, N. Buisine, L.M. Sachs, D.R. Buchholz, T. Kwon, H. Smith-Parker, M. Gridi-Papp, M.J. Ryan, R.D. Denton, J.H. Malone, J.B. Wallingford, A.F. Straight, R. Heald, D. Hockemeyer, R.M. Harland, and D.S. Rokhsar. 2021. Conserved chromatin and repetitive patterns reveal slow genome evolution in frogs. bioRxiv. 2021.10.18.464293.

9. Brown, K.S., M.D. Blower, T.J. Maresca, T.C. Grammer, R.M. Harland, and R. Heald. 2007. Xenopus tropicalis egg extracts provide insight into scaling of the mitotic spindle. J. Cell Biol. 176:765–70. doi:10.1083/jcb.200610043.

10. Bürki, E. 1985. The expression of creatine kinase isozymes in Xenopus tropicalis, Xenopus laevis laevis, and their viable hybrid. Biochem. Genet. 23:73–88. doi:10.1007/BF00499114.

11. Carroll, C.W., M.C.C. Silva, K.M. Godek, L.E.T. Jansen, and A.F. Straight. 2009. Centromere assembly requires the direct recognition of CENP-A nucleosomes by CENP-N. Nat. Cell Biol. 11:896–902. doi:10.1038/ncb1899.

12. Chittori, S., J. Hong, H. Saunders, H. Feng, R. Ghirlando, A.E. Kelly, Y. Bai, and S. Subramaniam. 2018. Structural mechanisms of centromeric nucleosome recognition by the kinetochore protein CENP-N. Science (80-.). 359:339–343. doi:10.1126/science.aar2781.

13. Colis, L., K. Peltonen, P. Sirajuddin, H. Liu, S. Sanders, G. Ernst, J.C. Barrow, and M. Laiho. 2014. DNA intercalator BMH-21 inhibits RNA polymerase I independent of DNA damage response. Oncotarget. 5:4361–4369. doi:10.18632/oncotarget.2020.

14. Deng, L., R.A. Wu, R. Sonneville, O. V. Kochenova, K. Labib, D. Pellman, and J.C. Walter. 2019. Mitotic CDK Promotes Replisome Disassembly, Fork Breakage, and Complex DNA Rearrangements. Mol. Cell. 73:915–929.e6. doi:10.1016/j.molcel.2018.12.021.

15. Dunleavy, E.M., D. Roche, H. Tagami, N. Lacoste, D. Ray-Gallet, Y. Nakamura, Y. Daigo, Y. Nakatani, and G. Almouzni-Pettinotti. 2009. HJURP Is a Cell-Cycle-Dependent Maintenance and Deposition Factor of CENP-A at Centromeres. Cell. 137:485–497. doi:10.1016/j.cell.2009.02.040.

16. Durica, D.S., and H.M. Krider. 1977. Studies on the ribosomal RNA cistrons in interspecific Drosophila hybrids. I. Nucleolar dominance. Dev. Biol. 59:62–74. doi:10.1016/0012-1606(77)90240-8.

17. Durkin, S.G., and T.W. Glover. 2007. Chromosome Fragile Sites. Annu. Rev. Genet. 41:169–192. doi:10.1146/annurev.genet.41.042007.165900.

18. Edelstein, A.D., M. a Tsuchida, N. Amodaj, H. Pinkard, R.D. Vale, and N. Stuurman. 2014. Advanced methods of microscope control using μManager software. J. Biol. Methods. 1:10. doi:10.14440/jbm.2014.36.

19. Edwards, N.S., and A.W. Murray. 2005. Identification of Xenopus CENP-A and an Associated Centromeric DNA Repeat. Mol. Biol. Cell. 16:1800–1810. doi:10.1091/mbc.E04.

20. Erhardt, S., B.G. Mellone, C.M. Betts, W. Zhang, G.H. Karpen, and A.F. Straight. 2008. Genome-wide analysis reveals a cell cycle-dependent mechanism controlling centromere propagation. J. Cell Biol. 183:805–818. doi:10.1083/jcb.200806038.

21. Falk, S.J., L.Y. Guo, N. Sekulic, E.M. Smoak, T. Mani, G.A. Logsdon, K. Gupta, L.E.T. Jansen, G.D. Van Duyne, S.A. Vinogradov, M.A. Lampson, and B.E. Black. 2015. CENP-C reshapes and stabilizes CENP-A nucleosomes at the centromere. Science (80-.). 348:699–704.

22. Foltz, D.R., L.E.T. Jansen, A.O. Bailey, J.R. Yates, E.A. Bassett, S. Wood, B.E. Black, and D.W. Cleveland. 2009. Centromere-Specific Assembly of CENP-A Nucleosomes Is Mediated by HJURP. Cell. 137:472–484. doi:10.1016/j.cell.2009.02.039.

23. French, B.T., and A.F. Straight. 2017. The Power of Xenopus Egg Extract for Reconstitution of Centromeres and Kinetochore Function. Prog Mol Subcell Biol. 56:59–84. doi:10.1007/978-3-319-58592-5.

24. French, B.T., F.G. Westhorpe, C. Limouse, and A.F. Straight. 2017. Xenopus laevis M18BP1 Directly Binds Existing CENP-A Nucleosomes to Promote Centromeric Chromatin Assembly. Dev. Cell. 42:190–199.e10. doi:10.1016/j.devcel.2017.06.021.

25. Fu, L., B. Niu, Z. Zhu, S. Wu, and W. Li. 2012. CD-HIT: Accelerated for clustering the next-generation sequencing data. Bioinformatics. 28:3150–3152. doi:10.1093/bioinformatics/bts565.

26. Fujiwara, A., S. Abe, E. Yamaha, F. Yamazaki, and M.C. Yoshida. 1997. Uniparental chromosome elimination in the early embryogenesis of the inviable salmonid hybrids between masu salmon female and rainbow trout male. Chromosoma. 106:44–52. doi:10.1007/s004120050223.

27. Gébrane-Younès, J., N. Fomproix, and D. Hernandez-Verdun. 1997. When rDNA transcription is arrested during mitosis, UBF is still associated with non-condensed rDNA. J. Cell Sci. 110:2429–2440.

28. Gernand, D., T. Rutten, A. Varshney, M. Rubtsova, S. Prodanovic, and C. Bru. 2005. Uniparental Chromosome Elimination at Mitosis and Interphase in Wheat and Pearl Millet Crosses Involves Micronucleus Formation, Progressive Heterochromatinization, and DNA Fragmentation. Plant Cell. 17:2431–2438. doi:10.1105/tpc.105.034249.).

29. Gibeaux, R., R. Acker, M. Kitaoka, G. Georgiou, I. van Kruijsbergen, B. Ford, E.M. Marcotte, D.K. Nomura, T. Kwon, G.J.C. Veenstra, and R. Heald. 2018. Paternal chromosome loss and metabolic crisis contribute to hybrid inviability in Xenopus. Nature. 553:337–341. doi:10.1038/nature25188.

30. Gibeaux, R., and R. Heald. 2019. Generation of Xenopus Haploid, Triploid, and Hybrid Embryos. Methods Mol. Biol. 1920:303–315. doi:10.1007/978-1-61779-210-6.

31. Gómez-González, B., and A. Aguilera. 2019. Transcription-mediated replication hindrance : a major driver of genome instability. Genes Dev. 33:1008–1026. doi:10.1101/gad.324517.119.Freely.

32. Grenfell, A.W., R. Heald, and M. Strzelecka. 2016. Mitotic noncoding RNA processing promotes kinetochore and spindle assembly in Xenopus. J. Cell Biol. 214:133–141. doi:10.1083/jcb.201604029.

33. Hannak, E., and R. Heald. 2006. Investigating mitotic spindle assembly and function in vitro using Xenopus laevis egg extracts. Nat. Protoc. 1:2305–2314. doi:10.1038/nprot.2006.396.

34. Henikoff, S., K. Ahmad, and H.S. Malik. 2011. The Centromere Paradox : Stable Inheritance with Rapidly Evolving DNA. Science (80-.). 293:1098–1102. doi:10.1126/science.1062939.

35. Hooff, J.J., E. Tromer, L.M. Wijk, B. Snel, and G.J. Kops. 2017. Evolutionary dynamics of the kinetochore network in eukaryotes as revealed by comparative genomics. EMBO Rep. 18:1559–1571. doi:10.15252/embr.201744102.

36. Hori, T., W.H. Shang, M. Hara, M. Ariyoshi, Y. Arimura, R. Fujita, H. Kurumizaka, and T. Fukagawa. 2017. Association of M18BP1/KNL2 with CENP-A Nucleosome Is Essential for Centromere Formation in Non-mammalian Vertebrates. Dev. Cell. 42:181–189.e3. doi:10.1016/j.devcel.2017.06.019.

37. Hu, H., Y. Liu, M. Wang, J. Fang, H. Huang, N. Yang, Y. Li, J. Wang, X. Yao, Y. Shi, G. Li, and R.M. Xu. 2011. Structure of a CENP-A-histone H4 heterodimer in complex with chaperone HJURP. Genes Dev. 25:901–906. doi:10.1101/gad.2045111.

38. Jagannathan, M., and Y.M. Yamashita. 2021. Defective Satellite DNA Clustering into Chromocenters Underlies Hybrid Incompatibility in Drosophila. Mol. Biol. Evol. 38:4977–4986. doi:10.1093/molbev/msab221.

39. Kabeche, L., H.D. Nguyen, R. Buisson, and L. Zou. 2018. A mitosis-specific and R loop-driven ATR pathway promotes faithful chromosome segregation SUPP. Science (80-.). 359:108–114. doi:10.1126/science.aan6490.

40. Kitaoka, M., R. Heald, and R. Gibeaux. 2018. Spindle assembly in egg extracts of the Marsabit clawed frog, Xenopus borealis. Cytoskeleton. 75:244–257. doi:10.1002/cm.21444.

41. Kumon, T., J. Ma, R.B. Akins, D. Stefanik, C.E. Nordgren, J. Kim, M.T. Levine, and M.A. Lampson. 2021. Parallel pathways for recruiting effector proteins determine centromere drive and suppression. Cell. 1–15. doi:10.1016/j.cell.2021.07.037.

42. Lee, H.Y., J.Y. Chou, L. Cheong, N.H. Chang, S.Y. Yang, and J.Y. Leu. 2008. Incompatibility of Nuclear and Mitochondrial Genomes Causes Hybrid Sterility between Two Yeast Species. Cell. 135:1065–1073. doi:10.1016/j.cell.2008.10.047.

43. Levy, D.L., and R. Heald. 2010. Nuclear Size Is Regulated by Importin A and Ntf2 in Xenopus. Cell. 143:288–298. doi:10.1016/j.cell.2010.09.012.

44. Lukacs, A., A.W. Thomae, P. Krueger, T. Schauer, A. V. Venkatasubramani, N.Y. Kochanova, W. Aftab, R. Choudhury, I. Forne, and A. Imhof. 2021. The integrity of the HMR complex is necessary for centromeric binding and reproductive isolation in Drosophila. PLoS Genet. 17:1–27. doi:10.1371/journal.pgen.1009744.

45. Ma, H., N. Marti Gutierrez, R. Morey, C. Van Dyken, E. Kang, T. Hayama, Y. Lee, Y. Li, R. Tippner-Hedges, D.P. Wolf, L.C. Laurent, and S. Mitalipov. 2016. Incompatibility between Nuclear and Mitochondrial Genomes Contributes to an Interspecies Reproductive Barrier. Cell Metab. 24:283–294. doi:10.1016/j.cmet.2016.06.012.

46. Maheshwari, S., and D.A. Barbash. 2011. The Genetics of Hybrid Incompatibilities. Annu. Rev. Genet. 45:331–355. doi:10.1146/annurev-genet-110410-132514.

47. Maheshwari, S., E.H. Tan, A. West, F.C.H. Franklin, L. Comai, and S.W.L. Chan. 2015. Naturally Occurring Differences in CENH3 Affect Chromosome Segregation in Zygotic Mitosis of Hybrids. PLoS Genet. 11:1–20. doi:10.1371/journal.pgen.1004970.

48. Malik, H.S., and S. Henikoff. 2001. Adaptive evolution of Cid, a centromere-specific histone in Drosophila. Genetics. 157:1293–1298. doi:10.1093/genetics/157.3.1293.

49. Malik, H.S., and S. Henikoff. 2009. Major Evolutionary Transitions in Centromere Complexity. Cell. 138:1067–1082. doi:10.1016/j.cell.2009.08.036.

50. Malik, H.S., D. Vermaak, and S. Henikoff. 2002. Recurrent evolution of DNA-binding motifs in the Drosophila centromeric histone. Proc. Natl. Acad. Sci. U. S. A. 99:1449–54. doi:10.1073/pnas.032664299.

51. Maresca, T.J., and R. Heald. 2006. Methods for studying spindle assembly and chromosome condensation in Xenopus egg extracts. Methods Mol. Biol. (Clifton, NJ). 322:459–474. doi:10.1007/978-1-59745-000-3_33.

52. Maric, M., T. Maculins, G. De Piccoli, and K. Labib. 2014. Cdc48 and a ubiquitin ligase drive disassembly of the CMG helicase at the end of DNA replication. Science (80-.). 346. doi:10.1126/science.1253596.

53. Milks, K.J., B. Moree, and A.F. Straight. 2009. Dissection of CENP-C – directed Centromere and Kinetochore Assembly. Mol. Biol. Cell. 20:4246–4255. doi:10.1091/mbc.E09.

54. Moree, B., C.B. Meyer, C.J. Fuller, and A.F. Straight. 2011. CENP-C recruits M18BP1 to centromeres to promote CENP-A chromatin assembly. J. Cell Biol. 194:855–871. doi:10.1083/jcb.201106079.

55. Narbonne, P., D.E. Simpson, and J.B. Gurdon. 2011. Deficient induction response in a Xenopus nucleocytoplasmic hybrid. PLoS Biol. 9:e1001197. doi:10.1371/journal.pbio.1001197.

56. Newport, J., and M. Kirschner. 1982. A major developmental transition in early xenopus embryos: I. characterization and timing of cellular changes at the midblastula stage. Cell. 30:675–686. doi:10.1016/0092-8674(82)90272-0.

57. Peltonen, K., L. Colis, H. Liu, R. Trivedi, M.S. Moubarek, H.M. Moore, B. Bai, M.A. Rudek, C.J. Bieberich, and M. Laiho. 2014. A targeting modality for destruction of RNA polymerase I that possesses anticancer activity. Cancer Cell. 25:77–90. doi:10.1016/j.ccr.2013.12.009.

58. Pentakota, S., K. Zhou, C. Smith, S. Maffini, A. Petrovic, G.P. Morgan, J.R. Weir, I.R. Vetter, A. Musacchio, and K. Luger. 2017. Decoding the centromeric nucleosome through CENP-N. Elife. 6:e33442. doi:10.7554/eLife.33442.

59. Pontremoli, C., D. Forni, U. Pozzoli, M. Clerici, R. Cagliani, and M. Sironi. 2021. Kinetochore proteins and microtubule-destabilizing factors are fast evolving in eutherian mammals. Mol. Ecol. 30:1505–1515. doi:10.1111/mec.15812.

60. De Robertis, E.M., and P. Black. 1979. Hybrids of Xenopus laevis and Xenopus borealis express proteins from both parents. Dev. Biol. 68:334–339. doi:10.1016/0012-1606(79)90267-7.

61. Rošić, S., F. Köhler, and S. Erhardt. 2014. Repetitive centromeric satellite RNA is essential for kinetochore formation and cell division. J. Cell Biol. 207:335–349. doi:10.1083/jcb.201404097.

62. Rosin, L., and B.G. Mellone. 2016. Co-evolving CENP-A and CAL1 Domains Mediate Centromeric CENP-A Deposition across Drosophila Species. Dev. Cell. 37:136–147. doi:10.1016/j.devcel.2016.03.021.

63. Rosin, L.F., and B.G. Mellone. 2017. Centromeres Drive a Hard Bargain. Trends Genet. 33:101–117. doi:10.1016/j.tig.2016.12.001.

64. Roure, V., B. Medina-Pritchard, V. Lazou, L. Rago, E. Anselm, D. Venegas, A.A. Jeyaprakash, and P. Heun. 2019. Reconstituting Drosophila Centromere Identity in Human Cells. Cell Rep. 29:464–479.e5. doi:10.1016/j.celrep.2019.08.067.

65. Roussel, P., C. André, L. Comai, and D. Hernandez-Verdun. 1996. The rDNA transcription machinery is assembled during mitosis in active NORs and absent in inactive NORs. J. Cell Biol. 133:235–246. doi:10.1083/jcb.133.2.235.

66. Sanei, M., R. Pickering, K. Kumke, S. Nasuda, and A. Houben. 2011. Loss of centromeric histone H3 (CENH3) from centromeres precedes uniparental chromosome elimination in interspecific barley hybirds. PNAS. 108:E498–E505. doi:10.1073/pnas.1103190108/-/DCSupplemental.www.pnas.org/cgi/doi/10.1073/pnas.1103190108.

67. Satyaki, P.R. V, T.N. Cuykendall, K.H.C. Wei, N.J. Brideau, H. Kwak, S. Aruna, P.M. Ferree, S. Ji, and D.A. Barbash. 2014. The Hmr and Lhr Hybrid Incompatibility Genes Suppress a Broad Range of Heterochromatic Repeats. PLoS Genet. 10. doi:10.1371/journal.pgen.1004240.

68. Schindelin, J., I. Arganda-Carreras, E. Frise, V. Kaynig, M. Longair, T. Pietzsch, S. Preibisch, C. Rueden, S. Saalfeld, B. Schmid, J.-Y. Tinevez, D.J. White, V. Hartenstein, K. Eliceiri, P. Tomancak, and A. Cardona. 2012. Fiji: an open-source platform for biological-image analysis. Nat. Methods. 9:676–682. doi:10.1038/nmeth.2019.

69. Session, A.M., Y. Uno, T. Kwon, J.A. Chapman, A. Toyoda, S. Takahashi, A. Fukui, A. Hikosaka, A. Suzuki, M. Kondo, S.J. van Heeringen, I. Quigley, S. Heinz, H. Ogino, H. Ochi, U. Hellsten, J.B. Lyons, O. Simakov, N. Putnam, J. Stites, Y. Kuroki, T. Tanaka, T. Michiue, M. Watanabe, O. Bogdanovic, R. Lister, G. Georgiou, S.S. Paranjpe, I. van Kruijsbergen, S. Shu, J. Carlson, T. Kinoshita, Y. Ohta, S. Mawaribuchi, J. Jenkins, J. Grimwood, J. Schmutz, T. Mitros, S. V. Mozaffari, Y. Suzuki, Y. Haramoto, T.S. Yamamoto, C. Takagi, R. Heald, K. Miller, C. Haudenschild, J. Kitzman, T. Nakayama, Y. Izutsu, J. Robert, J. Fortriede, K. Burns, V. Lotay, K. Karimi, Y. Yasuoka, D.S. Dichmann, M.F. Flajnik, D.W. Houston, J. Shendure, L. DuPasquier, P.D. Vize, A.M. Zorn, M. Ito, E.M. Marcotte, J.B. Wallingford, Y. Ito, M. Asashima, N. Ueno, Y. Matsuda, G.J.C. Veenstra, A. Fujiyama, R.M. Harland, M. Taira, and D.S. Rokhsar. 2016. Genome evolution in the allotetraploid frog Xenopus laevis. Nature. 538:336–343. doi:10.1038/nature19840.

70. Shiokawa, K., R. Kurashima, and J. Shinga. 1994. Temporal control of gene expression from endogenous and exogenously-introduced DNAs in early embryogenesis of Xenopus laevis. Int. J. Dev. Biol. 38:249–255. doi:10.1387/ijdb.7526881.

71. Shono, N., J.I. Ohzeki, K. Otake, N.M.C. Martins, T. Nagase, H. Kimura, V. Larionov, W.C. Earnshaw, and H. Masumoto. 2015. CENP-C and CENP-I are key connecting factors for kinetochore and CENP-A assembly. J. Cell Sci. 128:4572–4587. doi:10.1242/jcs.180786.

72. Smith, O.K., C. Limouse, K.A. Fryer, N.A. Teran, K. Sundararajan, R. Heald, and A.F. Straight. 2021. Identification and characterization of centromeric sequences in Xenopus laevis. Genome Res. 31:958–967. doi:10.1101/gr.267781.120.

73. Stellfox, M.E., A.O. Bailey, and D.R. Foltz. 2013. Putting CENP-A in its place. Cell. Mol. Life Sci. 70:387–406. doi:10.1007/s00018-012-1048-8.

74. Thomae, A.W., G.O.M. Schade, J. Padeken, M. Borath, I. Vetter, E. Kremmer, P. Heun, and A. Imhof. 2013. A Pair of Centromeric Proteins Mediates Reproductive Isolation in Drosophila Species. Dev. Cell. 27:412–424. doi:10.1016/j.devcel.2013.10.001.

75. Tian, T., X. Li, Y. Liu, C. Wang, X. Liu, G. Bi, X. Zhang, X. Yao, Z.H. Zhou, and J. Zang. 2018. Molecular basis for CENP-N recognition of CENP-A nucleosome on the human kinetochore. Cell Res. 28:374–378. doi:10.1038/cr.2018.13.

76. Woodland, H.R., and J.E.M. Ballantine. 1980. Paternal gene expression in developing hybrid embryos of Xenopus laevis and Xenopus borealis. J. Embryol. Exp. Morphol. 60:359–372.

